# Deep genome annotation of the opportunistic human pathogen *Streptococcus pneumoniae* D39

**DOI:** 10.1101/283663

**Authors:** Jelle Slager, Rieza Aprianto, Jan-Willem Veening

## Abstract

A precise understanding of the genomic organization into transcriptional units and their regulation is essential for our comprehension of opportunistic human pathogens and how they cause disease. Using single-molecule realtime (PacBio) sequencing we unambiguously determined the genome sequence of *Streptococcus pneumoniae* strain D39 and revealed several inversions previously undetected by short-read sequencing. Significantly, a chromosomal inversion results in antigenic variation of PhtD, an important surface-exposed virulence factor. We generated a new genome annotation using automated tools, followed by manual curation, reflecting the current knowledge in the field. By combining sequence-driven terminator prediction, deep paired-end transcriptome sequencing and enrichment of primary transcripts by Cappable-Seq, we mapped 1,015 transcriptional start sites and 748 termination sites. We show that the pneumococcal transcriptional landscape is complex and includes many secondary, antisense and internal promoters. Using this new genomic map, we identified several new small RNAs (sRNAs), RNA switches (including sixteen previously misidentified as sRNAs), and antisense RNAs. In total, we annotated 89 new protein-encoding genes, 34 sRNAs and 165 pseudogenes, bringing the *S. pneumoniae* D39 repertoire to 2,146 genetic elements. We report operon structures and observed that 9% of operons are leaderless. The genome data is accessible in an online resource called PneumoBrowse (https://veeninglab.com/pneumobrowsel) providing one of the most complete inventories of a bacterial genome to date. PneumoBrowse will accelerate pneumococcal research and the development of new prevention and treatment strategies.

## INTRODUCTION

Ceaseless technological advances have revolutionized our capability to determine genome sequences as well as our ability to identify and annotate functional elements, including transcriptional units on these genomes. Several resources have been developed to organize current knowledge on the important opportunistic human pathogen *Streptococcus pneumoniae*, or the pneumococcus (1–3). However, an accurate genome map with an up-to-date and extensively curated genome annotation, is missing.

The enormous increase of genomic data on various servers, such as NCBI and EBI, and the associated decrease in consistency has, in recent years, led to the Prokaryotic RefSeq Genome Re-annotation Project. Every bacterial genome present in the NCBI database was re-annotated using the so-called Prokaryotic Genome Annotation Pipeline (PGAP, (4)), with the goal of increasing the quality and consistency ofthe many available annotations. This Herculean effort indeed created a more consistent set of annotations that facilitates the propagation and interpolation of scientific findings in individual bacteria to general phenomena, valid in larger groups of organisms. On the other hand, a wealth of information is already available for well-studied bacteria like the pneumococcus. Therefore, a separate, manually curated annotation is essential to maintain oversight of the current knowledge in the field. Hence, we generated a resource for the pneumococcal research community that contains the most up-to-date information on the D39 genome, including its DNA sequence, transcript boundaries, operon structures and functional annotation. Notably, strain D39 is one of the workhorses in research on pneumococcal biology and pathogenesis. We analyzed the genome in detail, using a combination of several different sequencing techniques and a novel, generally applicable analysis pipeline (Figure 1).

**Figure 1.**
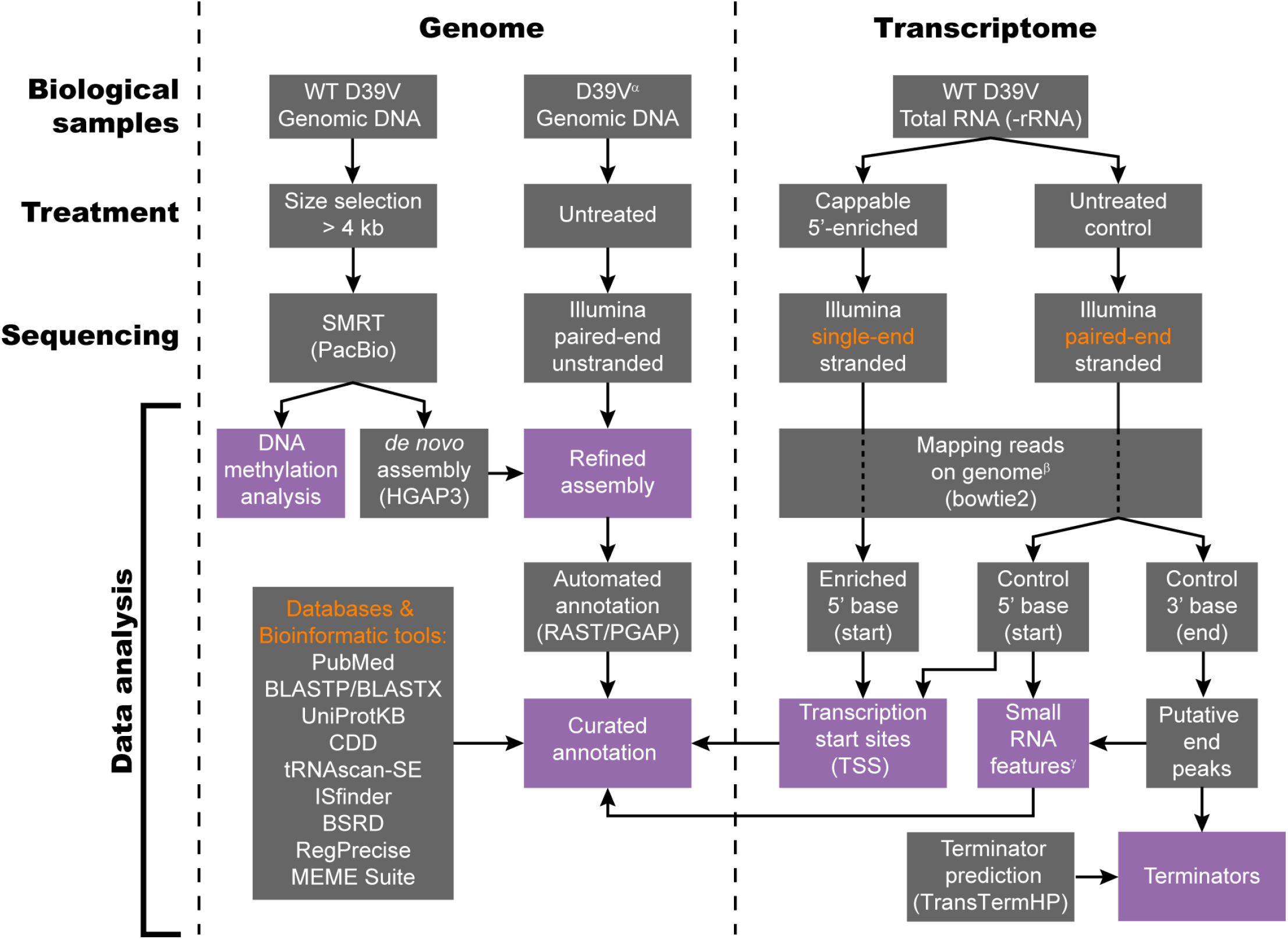
Data analysis pipeline used for genome assembly and annotation. Left. DNA level: the genome sequence of D39V was determined by SMRT sequencing, supported by previously published Illumina data (10, 25). Automated annotation by the RAST (13) and PGAP (4) annotation pipelines was followed by curation based on information from literature and a variety of databases and bioinformatic tools. Right. RNA level: Cappable-seq (7) was utilized to identify transcription start sites. Simultaneously, putative transcript ends were identified by combining reverse reads from paired-end, stranded sequencing of the control sample (i.e. not 5’-enriched). Terminators were annotated when such putative transcript ends overlapped with stem loops predicted by TransTermHP (22). Finally, local fragment size enrichment in the paired-end sequencing data was used to identify putative small RNA features. ^α^D39 derivative (*bgaA*::P_*ssbB*_-*luc*; GEO accessions GSE54199 and GSE69729). ^β^The first 1 kbp of the genome file was duplicated at the end, to allow mapping over FASTA boundaries. ^γ^Analysis was performed with only sequencing pairs that map uniquely to the genome.

Using Single Molecule Real-Time (SMRT, PacBio RS II) sequencing, we sequenced the genome of the stock of serotype 2 *S. pneumoniae* strain D39 in the Veening laboratory, hereafter referred to as strain D39V. This strain is a far descendant of the original Avery strain that was used to demonstrate that DNA is the carrier of hereditary information ((5, 6), **Supplementary Figure S1**). Combining Cappable-seq (7), a novel sRNA detection method and several bioinformatic annotation tools, we deeply annotated the pneumococcal genome and transcriptome.

Finally, we created PneumoBrowse, an intuitive and accessible genome browser (https://veeninglab.com/pneumobrowse), based on JBrowse (8). PneumoBrowse provides a graphical and user-friendly interface to explore the genomic and transcriptomic landscape of *S. pneumoniae* D39V and allows direct linking to gene expression and co-expression data in PneumoExpress (9). The reported annotation pipeline and accompanying genome browser provide one of the best curated bacterial genomes currently available and may facilitate rapid and accurate annotation of other bacterial genomes. We anticipate that PneumoBrowse will significantly accelerate the pneumococcal research field and hence speed-up the discovery of new drug targets and vaccine candidates for this devastating global opportunistic human pathogen.

## MATERIALS AND METHODS

### Culturing of *S. pneumoniae* D39 and strain construction

*S. pneumoniae* was routinely cultured without antibiotics. Transformation, strain construction and preparation of growth media are described in detail in the **Supplementary Methods**. Bacterial strains are listed in **Supplementary Table S1** and oligonucleotides in **Supplementary Table S2**.

### Growth, luciferase and GFP assays

Cells were routinely pre-cultured in C+Y medium (unless stated otherwise: pH 6.8, standing culture at ambient air) until an OD_600_ of 0.4, and then diluted 1:100 into fresh medium in a 96-wells plate. All assays were performed in a Tecan Infinite 200 PRO at 37°C. Luciferase assays were performed in C+Y with 0.25 mg·ml^−1^ D-luciferin sodium salt and signals were normalized by OD_595_. Fluorescence signals were normalized using data from a parental *gfp*-free strain. Growth assays of *lacD*-repaired strains were performed in C+Y with either 10.1 mM galactose or 10.1 mM glucose as main carbon source.

### DNA and RNA isolation, primary transcript enrichment and sequencing

*S. pneumoniae* chromosomal DNA was isolated as described previously (10). A 6/8 kbp insert library for SMRT sequencing, with a lower cut-off of 4kbp, was created by the Functional Genomics Center Zurich (FGCZ) and was then sequenced using a PacBio RS II machine.

D39V samples for RNA-seq were pre-cultured in suitable medium before inoculation (1:100) into four infection-relevant conditions, mimicking (i) lung, (ii) cerebral spinal fluid (CSF), or (iii) fever in CSF-like medium, and (iv) late competence (20 min after CSP addition) in C+Y medium. Composition of media, a detailed description of conditions and the total RNA isolation protocol are described in the accompanying paper by Aprianto et al. (9). Isolated RNA was sent to vertis Biotechnologie AG for sequencing. Total RNA from the four conditions was combined in an equimolar fashion and the pooled RNA was divided into two portions. The first portion was directly enriched for primary transcripts (Cappable-seq, (7)) and, after stranded library preparation according to the manufacturer’s recommendations, sequenced on Illumina NextSeq in single-end (SE) mode. The second RNA portion was rRNA-depleted, using the Ribo-Zero rRNA Removal Kit for Bacteria (Illumina), and sequenced on Illumina NextSeq in paired-end (PE) mode. RNA-seq data was mapped to the newly assembled genome using Bowtie 2 (11).

### *De novo* assembly of the D39V genome and DNA modification analysis

Analysis of SMRT sequencing data was performed with the SMRT tools analysis package (DNA Link, Inc., Seoul, Korea). *De novo* genome assembly was performed using the Hierarchical Genome Assembly Process (HGAP3) module of the PacBio SMRT portal version 2.3.0. This resulted in two contigs: one of over 2 Mbp with 250-500x coverage, and one of 12 kbp with 5-25x coverage. The latter, small contig was discarded based on its low coverage and high sequence similarity with a highly repetitive segment of the larger contig. The large contig was circularized manually and then rotated such that *dnaA* was positioned on the positive strand, starting on the first nucleotide. Previously published Illumina data of our D39 strain (GEO accessions GSE54199 and GSE69729) were mapped on the new assembly, using *breseq* (12), to identify potential discrepancies. Identified loci of potential mistakes in the assembly were verified by Sanger sequencing, leading to the correction of a single mistake. DNA modification analysis was performed using the ‘RS_Modification_and_Motif_Analysis.1’ module in the SMRT portal, with a QV cut-off of 100. A PCR-based assay to determine the absence or presence of plasmid pDP1 is described in **Supplementary Methods**.

### Automated and curated annotation

The assembled genome sequence was annotated automatically, using PGAP ((4), executed October 2015) and RAST ((13), executed June 2016). The results of both annotations were compared and for each discrepancy, support was searched. Among the support used were scientific publications (PubMed), highly similar features (BLAST, (14)), reviewed UniProtKB entries (15) and detected conserved domains as found by CD-Search (16). When no support was found for either the PGAP or RAST annotation, the latter was used. Similarly, annotations of conserved features in strain R6 (NC_003098.1) were adopted when sufficient evidence was available. Finally, an extensive literature search was performed, with locus tags and (if available) gene names from the old D39 annotation (prefix: ‘SPD_’) as query. When identical features were present in R6, a similar search was performed with R6 locus tags (prefix ‘spr’) and gene names. Using the resulting literature, the annotation was further refined. Duplicate gene names were also resolved during curation.

CDS pseudogenes were detected by performing a BLASTX search against the NCBI non-redundant protein database, using the DNA sequence of two neighboring genes and their intergenic region as query. If the full-length protein was found, the two (or more, after another BLASTX iteration) genes were merged into one pseudogene.

Furthermore, sRNAs and RNA switches, transcriptional start sites (TSSs) and terminators, transcription-regulatory sequences and other useful features (all described below) were added to the annotation. Finally, detected transcript borders (TSSs and terminators) were used to refine coordinates of annotated features (e.g. alternative translational initiation sites). Afterwards, the quality of genome-wide translational initiation site (TIS) calls was evaluated using ‘assess_TIS_annotation.py’ (17).

All publications used in the curation process are listed in the **Supplementary Data**.

Conveniently, RAST identified pneumococcus-specific repeat regions: BOX elements (18), Repeat Units of the Pneumococcus (RUPs, (19)) and *Streptococcus pneumoniae* Rho-Independent Termination Elements (SPRITEs, (20)), which we included in our D39V annotation. Additionally, ISfinder (21) was used to locate Insertion Sequences (IS elements).

### Normalized start and end counts and complete coverage of sequenced fragments

The start and (for paired-end data) end positions of sequenced fragments were extracted from the sequence alignment map (SAM) produced by Bowtie 2. The positions were used to build strand-specific, single-nucleotide resolution frequency tables (start counts, end counts and coverage). For paired-end data, coverage was calculated from the entire inferred fragment (i.e. including the region between mapping sites of mate reads). Start counts, end counts and coverage were each normalized by division by the summed genome-wide coverage, excluding positions within 30 nts of rRNA genes.

### Identification of transcriptional terminators

Putative Rho-independent terminator structures were predicted with TransTermHP (22), with a minimum confidence level of 60. Calling of ‘putative coverage termination peaks’ in the paired-end sequencing data of the control library is described in detail in **Supplementary Methods**. When such a peak was found to overlap with the 3’-poly(U)-tract of a predicted terminator, the combination of both elements was annotated as a high-confidence (HC) terminator. Terminator efficiency was determined by the total number of fragments ending in a coverage termination peak, as a percentage of all fragments covering the peak (i.e. including non-terminated fragments).

### Detection of small RNA features

For each putative coverage termination peak (see above), fragments from the SAM file that ended inside the peak region were extracted. Of those reads, a peak-specific fragment size distribution was built and compared to the library-wide fragment size distributions (see **Supplementary Methods**). A putative sRNA was defined by several criteria: (i) the termination efficiency of the coverage termination peak should be above 30% (see above for the definition of termination efficiency), (ii) the relative abundance of the predicted sRNA length should be more than 25-fold higher than the corresponding abundance in the library-wide distribution, (iii) the predicted sRNA should be completely covered at least 15x for HC terminators and at least 200x for non-HC terminators. The entire process was repeated once more, now also excluding all detected putative sRNAs from the library-wide size distribution. For the scope of this paper, only predicted sRNAs that did not significantly overlap already annotated features were considered.

A candidate sRNA was annotated (either as sRNA or RNA switch) when either (i) a matching entry, with a specified function, was found in RFAM (23) and/or BSRD (24) databases; (ii) the sRNA was validated by Northern blotting in previous studies; or (iii) at least two transcription-regulatory elements were detected (i.e. transcriptional start or termination sites, or sigma factor binding sites).

### Transcription start site identification

Normalized start counts from 5’-enriched and control libraries were compared. Importantly, normalization was performed excluding reads that mapped within 30 bps of rRNA genes. An initial list was built of unclustered TSSs, which have (i) at least 2.5-fold higher normalized start counts in the 5’-enriched library, compared to the control library, and (ii) a minimum normalized start count of 2 (corresponding to 29 reads) in the 5’-enriched library. Subsequently, TSS candidates closer than 10 nucleotides were clustered, conserving the candidate with the highest start count in the 5’-enriched library. Finally, if the 5’-enriched start count of a candidate TSS was exceeded by the value at the nucleotide immediately upstream, the latter was annotated as TSS instead. The remaining, clustered TSSs are referred to as high-confidence (HC) TSSs. To account for rapid dephosphorylation of transcripts, we included a set of 34 lower confidence (LC) TSSs in our annotation, which were not overrepresented in the 5’-enriched library, but that did meet a set of strict criteria: (i) normalized start count in the control library was above 10 (corresponding to 222 reads), (ii) a TATAAT motif (with a maximum of 1 mismatch) was present in the 5-15 nucleotides upstream, (iii) the nucleotide was not immediately downstream of a processed tRNA, and (iv) the nucleotide was in an intergenic region. If multiple LC-TSSs were predicted in one intergenic region, only the strongest one was annotated. If a HC-TSS was present in the same intergenic region, the LC-TSS was only annotated when its 5’-enriched start count exceeded that of the HC-TSS. TSS classification and prediction of regulatory motifs is described in **Supplementary Methods**.

### Operon prediction and leaderless transcripts

Defining an operon as a set of one or more genes controlled by a single promoter, putative operons were predicted for each primary TSS. Two consecutive features on the same strand were predicted to be in the same operon if (i) their expression across 22 infection-relevant conditions was strongly correlated (correlation value > 0.75 (9)) and (ii) no strong terminator (>80% efficient) was found between the features. In a total of 70 leaderless transcripts, the TSS was found to overlap with the translation initiation site of the first encoded feature in the operon.

### PneumoBrowse

Pneumobrowse (https://veeninglab.com/pneumobrowse: alternative: http://jbrowse.molgenrug.nl/pneumobrowse) is based on JBrowse (8), supplemented with plugins *jbrowse-dark-theme* and *SitewideNotices* (https://github.com/erasche), and *ScreenShotPlugin, HierarchicalCheckboxPlugin* and *StrandedPlotPlugin* (BioRxiv: https://doi.org/10.1101/212654). Annotated elements were divided over six annotation tracks: (i) genes (includes pseudogenes, shown in grey), (ii) putative operons, (iii) TSSs and terminators, (iv) Predicted regulatory features, including TSSs and terminators, (v) repeats, and (vi) other features. Additionally, full coverage tracks are available, along with start and end counts. A user manual as well as an update history of PneumoBrowse is available on the home page.

## RESULTS

### *De novo* assembly yields a single circular chromosome

We performed *de novo* genome assembly using SMRT sequencing data, followed by polishing with high-confidence Illumina reads, obtained in previous studies (10, 25). Since this data was derived from a derivative of D39, regions of potential discrepancy were investigated using Sanger sequencing. In the end, we needed to correct the SMRT assembly in only one location. The described approach yielded a single chromosomal sequence of 2,046,572 base pairs, which was deposited to GenBank (accession number CP027540).

### D39V did not suffer disruptive mutations compared to ancestral strain NCTC 7466

We then compared the newly assembled genome with the sequence previously established in the Winkler laboratory sequence of D39 (D39W, (6)), which is slightly closer to the original Avery isolate (**Supplementary Figure S1**). We observed similar sequences, but with some notable differences (Table 1, Figure 2A). Furthermore, we crosschecked both sequences with the genome sequence of the ancestral strain NCTC 7466 (ENA accession number ERS1022033), which was recently sequenced with SMRT technology, as part of the NCTC 3000 initiative (https://www.phe-culturecollections.org.uk/collections/nctc-3000-project.aspx). Interestingly, D39V matches NCTC 7466 in all gene-disruptive discrepancies (e.g. frameshifts and a chromosomal inversion, see below). Most of these sites are characterized by their repetitive nature (e.g. homopolymeric runs or long repeated sequences), which may serve as a source of biological variation and thereby explain observed differences with the D39W genome. Differences among D39V, D39W and the resequenced ancestral NCTC 7466 strain are confined to a limited number of SNPs, indels and regions of genetic inversion, with unknown consequences for pneumococcal fitness. None of these changes attenuate the virulence of these strains in animal models, and these polymorphisms emphasize the dynamic nature of the pneumococcal genome. Notably, there are two sites where both the D39W and D39V assemblies differ from the resequenced ancestral NCTC 7466 strain (Table 1, **Supplementary Figure S2U-V**). Firstly, the ancestral strain harbors a mutation in *rrlC* (SPV_1814), one of four copies of the gene encoding 23S ribosomal RNA. It is not clear if this is a technical artefact in one of the assemblies (due to the large repeat size in this region), or an actual biological difference. Secondly, we observed a mutation in the upstream region of *cbpM* (SPV_1248) in both D39W and D39V.

**Figure 2.**
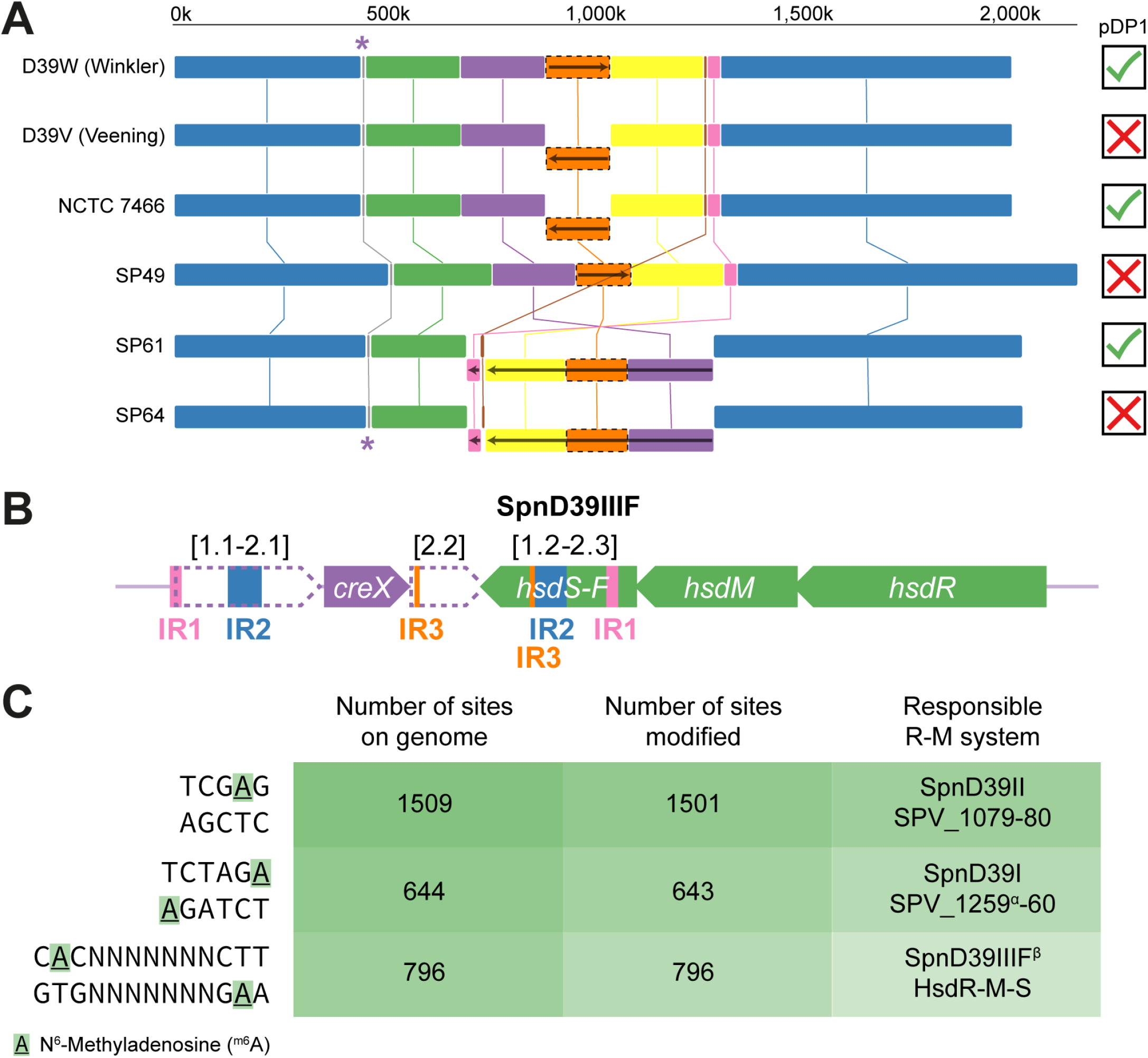
Multiple genome alignment. **A**. Multiple genome sequence alignment of D39W, D39V, NCTC 7466, and clinical isolates SP49, SP61, and SP64 (33) reveals multiple ter-symmetrical chromosomal inversions. Identical colors indicate similar sequences, while blocks shown below the main genome level and carrying a reverse arrow signify inverted sequences relative to the D39W assembly. The absence/presence of the pDP1 (or similar) plasmid is indicated with a cross/checkmark. Asterisks indicate the position of the *hsdS* locus. **B**. Genomic layout of the *hsdS* region. As reported by Manso et al. (28), the region contains three sets of inverted repeats (IR1-3), that are used by CreX to reorganize the locus. Thereby, six different variants (A-F) of methyltransferase specificity subunit HsdS can be generated, each leading to a distinct methylation motif. SMRT sequencing of D39V revealed that the locus exists predominantly in the F-configuration, consisting of N-terminal variant 2 (i.e. 1.2) and C-terminal variant 3 (i.e. 2.3). **C**. Motifs that were detected to be specifically modified in D39V SMRT data (see **Supplementary Methods**). Manso et al. identified the same motifs and reported the responsible methyltransferases. ^α^SPV_1259 (encoding the R-M system endonuclease) is a pseudogene, due to a nonsense mutation. ^β^The observed C<u>**A**</u>C-N_7_-CTT motif perfectly matches the HsdS-F motif predicted by Manso et al.

**Table 1.** Differences between old and new genome assembly. The genomic sequences of the old (D39W, CP000410.1) and new (D39V, CP027540) genome assemblies were compared, revealing 14 SNPs, 3 insertions, and 2 deletions. Additionally, a repeat expansion in *pavB*, several rearrangements in the *hsdS* locus and, most strikingly, a 162 kbp (8% of the genome) chromosomal inversion were observed. Finally, both sequences were compared to the recently released PacBio sequence of ancestral strain NCTC 7466 (ENA accession number ERS1022033). For each observed difference between the D39W and D39V assemblies, the variant matching the ancestral strain is displayed in boldface. ^*α*^Locus falls within the inverted *ter* region and the forward strain in the new assembly is therefore the reverse complementary of the old sequence (CP000410.1). ^β^Region is part of a larger pseudogene in the new annotation. ^γ^Only found in one of two D39 stocks in our laboratory. ^δ^Sequencing errors were identified in two locations of the D39W assembly, which, after recent correction, match the D39V assembly: a T was added at position 297,022, shifting SPD_0299 and SPD_0300 into the same reading frame (now annotated as SPD_0300/SPV_2141) and base 303,240 was updated from A to G (resulting in PBP2X N311D). ^ε^These reported changes are relative to NCTC 7466.

| D39W<sup>5</sup><br>position | D39V<br>position | Locus | Change | Consequence | Note |
| --- | --- | --- | --- | --- | --- |
| 80663-83343 | **80663-83799** | SPD_0080 (*pavB*) | Repeat expansion (6x>7x) | Repeat expansion in PavB (6x>7x) |  |
| **174318** | 174774 | SPD_0170 (*ruvA*) | G>A | RuvA V52V (GTG>GTA) |  |
| 458088-462242 | 458545-462699 | SPD_0450-55 (*hsdRMS*, *creX*) | Multiple rearrangements | HsdS type A>F | (Manso et al. 2014) |
| **462212** | 458575 | SPD_0453 (*hsdS*) | A>G | Imperfect > perfect inverted repeat | Inside rearranged region |
| **675950** | 676407 | SPD_0657 → / → SPD_0658 (*prfB*) | C>A | Intergenic (+163/-51 nt) | In 5' UTR of *prfB* |
| **775672** | 776129 | SPD_0764 (*sufS*) | G>A | SufS G318R (GGA>AGA) |  |
| 816157 | **816615** | SPD_0800<sup>8</sup> | +G | Frameshift (347/360 nt) |  |
| 901217-1062944 | **901675-1063403** | SPD_0889-1037 | Inversion | Swap of 3' ends of *phtB* (SPD_1037) and *phtD* (SPD_0889) |  |
| **934443** | 1030177 | SPD_0921 (*ccrB*) | A>G<sup>a</sup> | CcrB Q286R (CAG>CGG) |  |
| 951536 | **1013083**<sup>v</sup> | SPD_0942 | +C<sup>a</sup> | Frameshift (198/783 nt) |  |
| **1035166** | 929453 | SPD_1016 (*rexA*) | C>A<sup>a</sup> | RexA A961D (GCT>GAT) |  |
| 1080119 | **1080577** | SPD_1050 (*lacD*) | ΔT | Frameshift (159/981 nt) |  |
| 1171761 | **1172219** | SPD_1137 | C>G | H431Q (CAC>CAG) |  |
| 1256813 | **1257270** | SPD_1224 (*budA*) ← / → SPD_1225 | ΔA | Intergenic (-100/-42 nt) |  |
| **1256937** | 1257394<sup>v</sup> | SPD_1225 | G>T | R28L (CGC>CTC) |  |
| 1672084 | **1672541** | SPD_1660 (*rdgB*) | G>A | RdgB T117I (ACA>ATA) |  |
| **1676516** | 1676973 | SPD_1664 (*treP*) | C>T | TreP G359D (GGC>GAC) |  |
| **1787708** | 1788165 | SPD_1793 | C>T | A2V (GCA>GTA) |  |
| **1977728** | 1978185 | SPD_2002 (*dltD*) | C>A | DltD V252F (GTC>TTC) |  |
| **2022372** | 2022829 | SPD_2045 (*mreC*) | A>G | MreC S186P (TCT>CCT) |  |

**Table 1.** Differences between old and new genome assembly.
|  |  |  |  |  |  |
| --- | --- | --- | --- | --- | --- |
| 1276806 | 1277263 | SPD_1248 (*cbpM*) ← / ← SPD_1249 | A>G<sup>ε</sup> | Intergenic (-116/+1338 nt) | Identical in D39W and D39V |
| 1799597 | 1800054 | SPD_1814 (*rrlC*) | T>C<sup>ε</sup> | *rrlC* different from other 3 *rrl* copies in NCTC 7466 (872/2904 nt) | Identical in D39W and D39V |

### Several SNPs and indel mutations observed in D39V assembly

Thirteen single nucleotide polymorphisms (SNPs) were detected upon comparison of D39W and D39V assemblies. Comparison with NCTC 7466 showed that eleven of these SNPs seem to reflect mutations in D39V, while D39W differs from NCTC 7466 in the other two cases (Table 1). One of the SNPs results in a silent mutation in the gene encoding RuvA, the Holliday junction DNA helicase, while another SNP was located in the 5’-untranslated region (5’-UTR) of *prfB*, encoding peptide chain release factor 2. The other eleven SNPs caused amino acid changes in various proteins, including cell shape-determining protein MreC. It should be noted that one of these SNPs, leading to an arginine to leucine change in the protein encoded by SPV_1225 (previously SPD_1225), was not found in an alternative D39 stock from our lab (**Supplementary Figure S1**). The same applies to an insertion of a cytosine causing a frameshift in SPV_0942 (previously SPD_0942; **Supplementary Figure S2J**). All other differences found, however, were identified in both of our stocks and are therefore likely to be more widespread. Among these differences are three more indel mutations (insertions or deletions), the genetic context and consequences of which are shown in **Supplementary Figure S2**. One of the indels is located in the promoter region of two diverging operons, with unknown consequences for gene expression (**Supplementary Figure S2N**). Secondly, we found an insertion in the region corresponding to SPD_0800 (D39W annotation). Here, we report this gene to be part of a pseudogene (annotated as SPV_2242) together with SPD_0801. Hence, the insertion probably is of little consequence. Finally, a deletion was observed in the beginning of *lacD*, encoding an important enzyme in the D-tagatose-6-phosphate pathway, relevant in galactose metabolism. The consequential absence of functional LacD may explain why the inactivation of the alternative Leloir pathway in D39 significantly hampered growth on galactose (26). We repaired *lacD* in D39V and, as expected, observed restored growth on galactose (Supplementary Figure S3). Interestingly, D39W was confirmed to indeed have an intact *lacD* gene (M. Winkler, personal communication), even though the resequenced ancestral NCTC 7466 contained the non-functional, frameshifted *lacD.* Since *lacD* was reported to be essential only in a small subset of studied conditions (27), differences in culturing conditions between the different laboratories may explain the selection of cells with an intact *lacD* gene.

### Varying repeat frequency in surface-exposed protein PavB

Pneumococcal adherence and virulence factor B (PavB) is encoded by SPV_0080. Our assembly shows that this gene contains a series of seven imperfect repeats of 450-456 bps in size. Interestingly, SPD_0080 in D39W contains only six of these repeats. If identical repeat units are indicated with an identical letter, the repeat region in SPV_0080 of D39V can be written as *ABBCBDE*, where *E* is truncated after 408 bps. Using the same letter code, SPD_0080 of D39W contains *ABBCDE*, thus lacking the third repeat of element B, which is isolated from the other copies in SPV_0080. Because D39V and NCTC 7466 contain the full-length version of the gene, we hypothesize that D39W lost one of the repeats, making the encoded protein 152 residues shorter.

### Configuration of variable *hsdS* region matches observed methylation pattern

A local rearrangement is found in the pneumococcal *hsdS* locus, encoding a three-component restriction-modification system (HsdRMS). Recombinase CreX facilitates local recombination, using three sets of inverted repeats, and can thereby rapidly rearrange the region into six possible configurations (SpnD39IIIA-F). This process results in six different versions of methyltransferase specificity subunit HsdS, each with its own sequence specificity and transcriptomic consequences (28, 29). The region is annotated in the A-configuration in D39W, while the F-configuration is predominant in D39V (Figure 2B). Moreover, we exploited methylation data, intrinsically present in SMRT data (29–31), and observed an enriched methylation motif that exactly matches the putative SpnD39IIIF motif predicted by Manso et al. (Figure 2C, **Supplementary Table S3**). All 796 genome sites matching this motif were found to be modified on both strands in 80-100% of all sequenced molecules (p < 10^−20^), stressing the power of SMRT sequencing in methylome analysis.

### A large chromosomal inversion occurred multiple times in pneumococcal evolution

We also observed a striking difference between D39V and D39W: a 162 kbp region containing the replication terminus was completely inverted (Figures 2A and 3), with D39V matching the configuration of the resequenced ancestral strain NCTC 7466. The inverted region is bordered by two inverted repeats of 1.3 kb in length. We noticed that the *xerS/ dif_SL_* site, responsible for chromosome dimer resolution and typically located directly opposite the origin of replication (32), is asymmetrically situated on the right replichore in D39V (Figure 3A), while the locus is much closer to the halfway point of the chromosome in the D39W assembly, suggesting that this configuration is the original one and the observed inversion in D39V and NCTC 7466 is a true genomic change. To confirm this, we performed a PCR-based assay, in which the two possible configurations yield different product sizes. Indeed, the results showed that two possible configurations of the region exist in different pneumococcal strains; multiple D39 stocks, TIGR4, BHN100 and PMEN-14 have matching terminus regions, while the opposite configuration was found in R6, Rx1, PMEN-2 and PMEN-18. We repeated the analysis for a set of seven and a set of five strains, each related by a series of sequential transformation events. All strains had the same *ter* orientation (*not shown*), suggesting that the inversion is relatively rare, even in competent cells. However, both configurations are found in various branches of the pneumococcal phylogenetic tree, indicating multiple incidences of this chromosomal inversion. Interestingly, a similar, even larger inversion was observed in two out of three recently-sequenced clinical isolates of *S. pneumoniae* (33) (Figure 2A), suggesting a larger role for chromosomal inversions in pneumococcal evolution.

**Figure 3.**
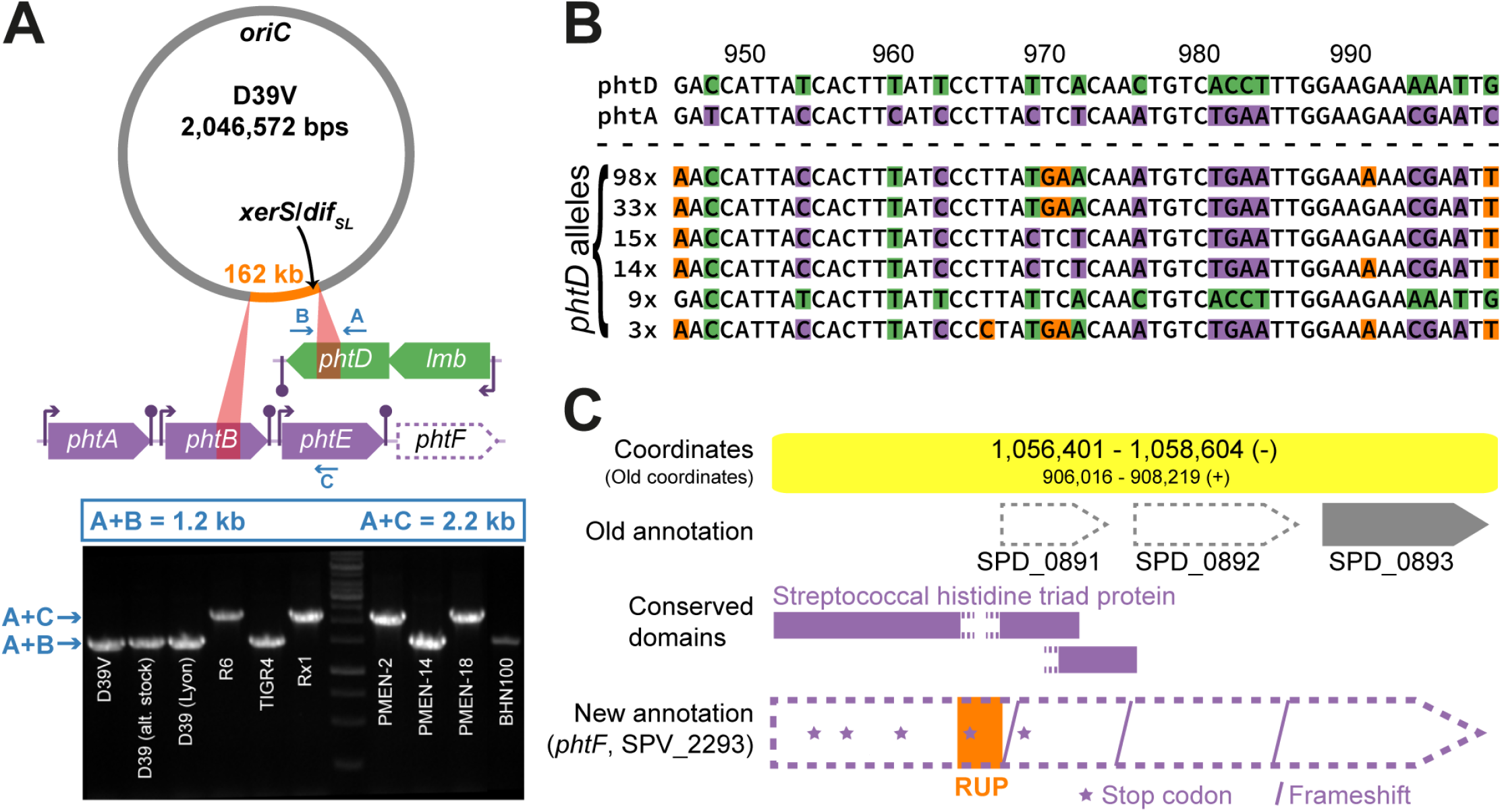
A large chromosomal inversion unveils antigenic variation of pneumococcal histidine triad proteins. **A**. Top: chromosomal location of the inverted 162 kb region (orange). Red triangles connect the location of the 1 kb inverted repeats bordering the inverted region and a zoom of the genetic context of the border areas, also showing that the inverted repeats are localized in the middle of genes *phtB* and *phtD*. Arrows marked with A, B and C indicate the target regions of oligonucleotides used in PCR analysis of the region. Bottom: PCR analysis of several pneumococcal strains (including both our D39 stocks and a stock from the Grangeasse lab, Lyon) shows that the inversion is a true phenomenon, rather than a technical artefact. PCR reactions are performed with all three primers present, such that the observed product size reports on the chromosomal configuration. **B**. A fragment of a Clustal Omega multiple sequence alignment of 172 reported *phtD* alleles (37) and D39V genes *phtD* and *phtA* exemplifies the dynamic nature of the genes encoding pneumococcal histidine triad proteins. Bases highlighted in green and purple match D39V *phtD* and *phtA*, respectively. Orange indicates that a base is different from both D39V genes, while white bases are identical in all sequences. **C**. Newly identified pseudogene, containing a RUP insertion and several frameshifts and nonsense mutations, that originally encoded a fifth pneumococcal histidine triad protein, and which we named *phtF.* Old (D39W) and new annotation (D39V) are shown, along with conserved domains predicted by CD-Search (16).

### Antigenic variation of histidine triad protein PhtD

Surprisingly, the repeat regions bordering the chromosomal inversion are located in the middle of *phtB* and *phtD* (Figure 3A), leading to an exchange of the C-terminal parts of their respective products, PhtB and PhtD. These are two out of four pneumococcal histidine triad (Pht) proteins, which are surface-exposed, interact with human host cells and are considered to be good vaccine candidates (34). In fact, PhtD was already used in several phase I/II clinical trials (e.g. (35, 36)). Yun et al. analyzed the diversity of *phtD* alleles from 172 clinical isolates and concluded that the sequence variation was minimal (37). However, this conclusion was biased by the fact that inverted chromosomes would not produce a PCR product in their set-up and a swap between PhtB and PhtD would remain undetected. Moreover, after detailed inspection of the mutations in the *phtD* alleles and comparison to other genes encoding Pht proteins (*phtA, phtB* and *phtE*), we found that many of the SNPs could be explained by recombination events between these genes, rather than by random mutation. For example, extensive exchange was seen between D39V *phtA* and *phtD*, as many of the aforementioned *phtD* alleles locally resembled phtA_D39V_ more closely than phtD_D39V_ (as exemplified by Figure 3B). Apparently, the repetitive nature of these genes allows for intragenomic recombination, causing *phtD* to become mosaic, rather than well-conserved. Finally, immediately downstream of *phtE* (Figure 3A), we identified a pseudogene (Figure 3C) that originally encoded a fifth histidine triad protein and which we named *phtF*, as previously suggested by Rioux et al. (38). The gene is disrupted by an inserted RUP element (see below) and several frameshifts and nonsense mutations, and therefore does not produce a functional protein. Nevertheless, *phtF* might still be relevant as a source of genetic diversity. Importantly, the variation in the DNA sequence of *phtD* is also propagated to the protein level. A 20 amino acid peptide (PhtD-pep19) was reported to have the highest reactivity with serum from patients suffering from invasive pneumococcal disease (39), rendering it a region of interest for vaccine development. A multiple sequence alignment of the 172 *phtD* alleles and the 5 *pht* genes from D39V revealed that this protein segment can potentially be varied in at least 8 out of 20 positions (**Supplementary Figure S4**). Taken together, these findings raise caution on the use of PhtD as a vaccine target.

### RNA-seq data and PCR analysis show loss of cryptic plasmid from strain D39V

Since SMRT technology is known to miss small plasmids in the assembly pipeline, we performed a PCR-based assay to check the presence of the cryptic pDP1 plasmid, reported in D39W (6, 40). To our surprise, the plasmid is absent in D39V, while clearly present in the ancestral NCTC 7466, as confirmed by a PCR-based assay (**Supplementary Figure S5**). Intriguingly, a BLASTN search suggested that *S. pneumoniae* Taiwan19F-14 (PMEN-14, CP000921), among other strains, integrated a degenerate version of the plasmid into its chromosome. Indeed, the PCR assay showed positive results for this strain. Additionally, we selected publicly available D39 RNA-seq datasets and mapped the sequencing reads specifically to the pDP1 reference sequence (Accession AF047696). The successful mapping of a significant number of reads indicated the presence of the plasmid in strains used in several studies ((41), SRX2613845; (42), SRX1725406; (28), SRX472966). In contrast, RNA-seq data of D39V (9–10, 25) contained zero reads that mapped to the plasmid, providing conclusive evidence that strain D39V lost the plasmid at some stage (**Supplementary Figure S1**). Similarly, based on Illumina DNA-seq data, we determined that of the three clinical isolates shown in Figure 2A, only SP61 contained a similar plasmid (33).

### Automation and manual curation yield up-to-date pneumococcal functional annotation

An initial annotation of the newly assembled D39V genome was produced by combining output from the RAST annotation engine (13) and the NCBI prokaryotic genome annotation pipeline (PGAP, (4)). We, then, proceeded with exhaustive manual curation to produce the final genome annotation (see **Materials and Methods** for details). All annotated CDS features without an equivalent feature in the D39W annotation or with updated coordinates are listed in **Supplementary Table S4**. Examples of the integration of recent research into the final annotation include cell division protein MapZ (43, 44), pleiotropic RNA-binding proteins KhpA and KhpB/EloR (45, 46), cell elongation protein CozE (47) and endolytic transglycosylase MltG (45, 48).

Additionally, we used tRNAscan-SE (49) to differentiate the four encoded tRNAs with a CAU anticodon into three categories (**Supplementary Table S5**): tRNAs used in either (i) translation initiation or (ii) elongation and (iii) the post-transcriptionally modified tRNA-Ile2, which decodes the AUA isoleucine codon (50).

Next, using BLASTX ((14), **Materials and Methods**), we identified and annotated 165 pseudogenes (**Supplementary Table S6**), two-fold more than reported previously (6). These non-functional transcriptional units may be the result of the insertion of repeat regions, nonsense and/or frameshift mutations and/or chromosomal rearrangements. Notably, 71 of 165 pseudogenes were found on IS elements (21), which are known to sometimes utilize alternative coding strategies, including programmed ribosomal slippage, producing a functional protein from an apparent pseudogene. Finally, we annotated 127 BOX elements (18), 106 RUPs (19), 29 SPRITEs (20) and 58 IS elements (21).

### RNA-seq coverage and transcription start site data allow improvement of annotated feature boundaries

Besides functional annotation, we also corrected the genomic coordinates of several features. First, we updated tRNA and rRNA boundaries (**Supplementary Table S5**), aided by RNA-seq coverage plots that were built from deduced paired-end sequenced fragments, rather than from just the sequencing reads. Most strikingly, we discovered that the original annotation of genes encoding 16S ribosomal RNA (*rrsA-D*) excluded the sequence required for ribosome binding site (RBS) recognition (51). Fortunately, neither RAST or PGAP reproduced this erroneous annotation and the D39V annotation includes these sites. Subsequently, we continued with correcting annotated translational initiation sites (TISs, start codons). While accurate TIS identification is challenging, 45 incorrectly annotated start codons could be identified by looking at the relative position of the corresponding transcriptional start sites (TSS/+1, described below). These TISs were corrected in the D39V annotation (**Supplementary Table S4**). Finally, we evaluated the genome-wide quality of TISs using a statistical model that compares the observed and expected distribution of the positions of alternative TISs relative to an annotated TIS (17). The developers suggested that a correlation score below 0.9 is indicative of poorly annotated TISs. In contrast to the D39W (0.899) and PGAP (0.873) annotations, our curated D39V annotation (0.945) excels on the test, emphasizing our annotation’s added value to pneumococcal research.

### Paired-end sequencing data contain the key to detection of small RNA features

After the sequence- and database-driven annotation process, we proceeded to study the transcriptome of *S. pneumoniae.* To maximize the number of expressed genes, we pooled RNA from cells grown at four different conditions (mimicking (i) lung, (ii) cerebral spinal fluid (CSF), or (iii) fever in CSF-like medium, and (iv) late competence (20 min after CSP addition) in C+Y medium; see (9)). Strand-specific, paired-end RNA-seq data of the control library was used to extract start and end points and fragment sizes of the sequenced fragments. In Figure 4A, the fragment size distribution of the entire library is shown, with a mode of approximately 150 nucleotides and a skew towards larger fragments. We applied a peak-calling routine to determine the putative 3’-ends of sequenced transcripts. For each of the identified peaks, we extracted all read pairs that were terminated in that specific peak region and compared the size distribution of that subset of sequenced fragments to the library-wide distribution to identify putative sRNAs (See **Materials and Methods** for more details). We focused on sRNA candidates that were found in intergenic regions. Using the combination of sequencing-driven detection, reported Northern blots, convincing homology with previously validated sRNAs, and/or presence of two or more regulatory features (e.g. TSSs and terminators, see below), we identified 63 small RNA features. We annotated 34 of these as sRNAs (Table 2) and 29 as RNA switches (**Supplementary Table S7**). While we may have missed sRNAs not expressed in the conditions studied here, inspection of our data in previously reported candidate regions (52–55) suggests that the utilized approaches in those studies result in a high false-positive rate.

**Figure 4.**
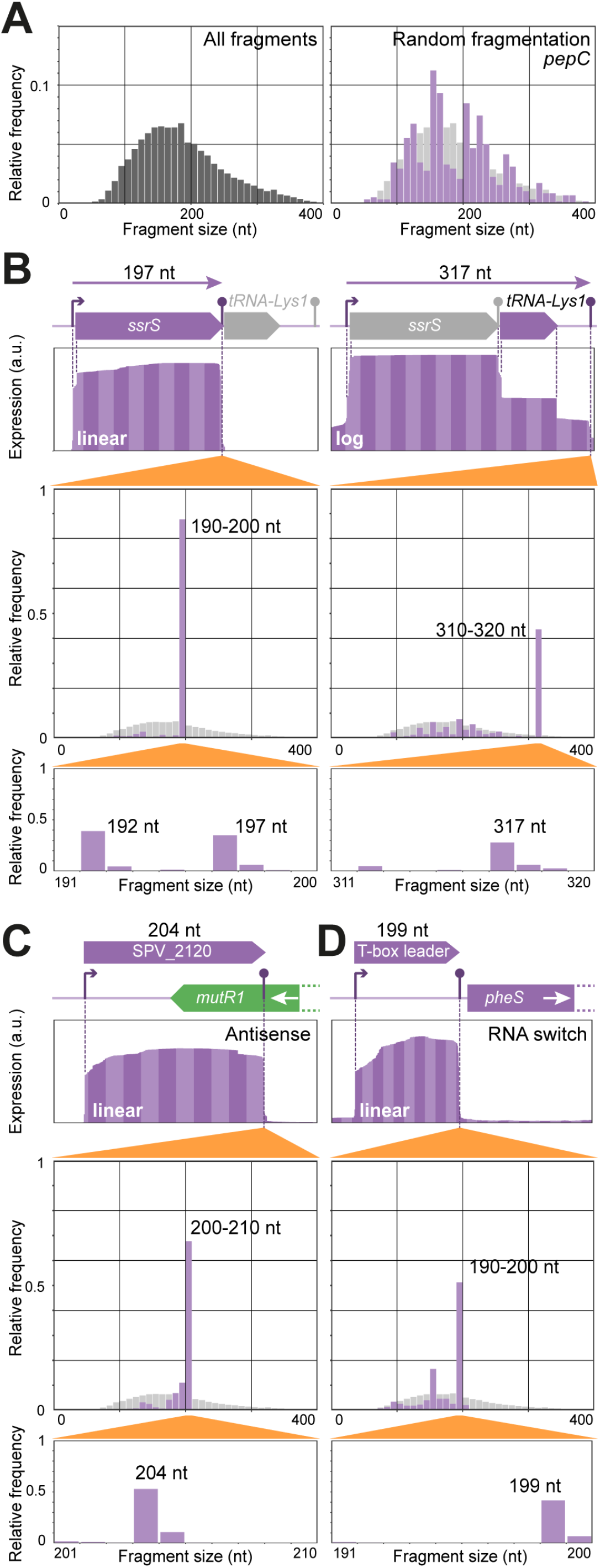
Detection of small RNAfeatures. **A**. Size distributions of entire sequencing library (left) and of fragments ending in the terminator region of *pepC* (right), as determined from paired-end sequencing. The negative control for sRNA detection, *pepC*, is much longer (1.3 kbp) than the typical fragment length in Illumina sequencing (100-350 bp). The plot is based on all sequenced molecules of which the 3’-end falls inside the downstream terminator region. The position of the 5’-end of each of these molecules is determined by random fragmentation in the library preparation. Therefore, its size distribution is expected to be comparable to that of the entire library (left plot and faint grey in right plot). **B-D**. Top in each panel: RNA-seq coverage plots, as calculated from paired-end sequenced fragments in the unprocessed control library (See **Supplementary Methods**). Bottom in each panel: size distributions (bin sizes 10 and 1) of fragments ending in the indicated terminator regions. **B**. Detection of ssrS (left) and joint *ssrS/tRNA-Lys1* (right) transcripts. Due to high abundance of *ssrS*, coverage is shown on log-scale in the right panel. Size distributions reveal 5’-processed and unprocessed *ssrS* (left) and full-length *ssrs*/*tRNA-Lys1* transcripts (right). **C**. Detection of an sRNA antisense to the 3’-region of *mutR1.* **D**. Detection of a T-box RNA switch structure upstream of *pheS*.

**Table 2.** All annotated small RNA features. Coordinates shown in boldface represent small RNA features not previously reported in *S. pneumoniae.* ^α^Not detected in this study, exact coordinates uncertain. ^β^Alternative terminator present. ^γ^Studies containing Northern blot validation are highlighted in boldface.

| D39V coordinates | Locus | Old locus | Gene name | Product | Note | Literature<sup>†</sup> |
| --- | --- | --- | --- | --- | --- | --- |
| 23967-24065 (+) | SPV_2078 |  | *ccnC* | Small regulatory RNA csRNA3 |  | 53, 54, **56** |
| 24632-24707 (+) | SPV_0026 | SPD_0026 | *scRNA* | scRNA | RNA component of signal recognition particle | 112 |
| **29658-29873 (+)** | SPV_2081 |  | *srf-01* | ncRNA of unknown function |  |  |
| 39980-40186 (+) | SPV_2084 |  | *srf-02* | ncRNA of unknown function |  | 53, **55** |
| **41719-41886 (+)** | SPV_2086 |  | *srf-03* | ncRNA of unknown function | Contains BOX element | 18, 114 |
| 132229-132308 (+) | SPV_2107 |  | *srf-04* | ncRNA of unknown function |  | 52, **57** |
| 149645-149877 (+)<sup>‡</sup> | SPV_2119 |  | *srf-05* | ncRNA of unknown function |  | 52, **57** |
| **150711-150914 (-)** | SPV_2120 |  | *srf-06* | ncRNA of unknown function | Antisense to *mutR1* (SPV_0144) |  |
| 212734-212881 (+) | SPV_2125 |  | *ccnE* | Small regulatory RNA csRNA5 | Involved in stationary phase autolysis | 53, 54, **55, 56** |
| 231599-231691 (+) | SPV_2129 |  | *ccnA* | Small regulatory RNA csRNA1 |  | 52, 53, 55, **56, 57** |
| 231786-231883 (+) | SPV_2130 |  | *ccnB* | Small regulatory RNA csRNA2 |  | 53, **56** |
| 232279-232354 (+)<sup>‡</sup> | SPV_2131 |  | *srf-07* | Type I addiction module antitoxin, Fst family |  | 55, 58 |
| 234171-234264 (+) | SPV_2133 |  | *ccnD* | Small regulatory RNA csRNA4 | Involved in stationary phase autolysis | 53, **56** |
| 282194-282479 (+) | SPV_2139 |  | *srf-08* | ncRNA of unknown function |  | 54, **55** |
| 344607-345007 (+) | SPV_0340 | SPD_0340 | *rnpB* | Ribonuclease P RNA component |  | 115 |
| 508697-508842 (+) | SPV_2185 |  | *srf-10* | ncRNA of unknown function |  | 55 |
| 587896-587989 (+) | SPV_2200 |  | *srf-11* | ncRNA of unknown function |  | 53, 54, **55** |
| 742022-742141 (-)<sup>‡</sup> | SPV_2226 |  | *srf-12* | ncRNA of unknown function | Implicated in competence | **54** |
| 781595-781939 (+) | SPV_0769 | SPD_0769 | *ssrA* | tmRNA |  | 53, **54**, 55, 112 |
| 826260-826587 (+) | SPV_2247 |  | *srf-13* | ncRNA of unknown function |  | 53, 54, 55 |
| 1037649-1037755 (-) | SPV_2291 |  | *srf-16* | ncRNA of unknown function |  | 55 |
| 1051910-1052049 (-) | SPV_2292 |  | *srf-17* | *asd* RNA motif |  | 52, 54, 55, **57**, 116 |
| **1079561-1079658 (-)** | SPV_2300 |  | *srf-18* | ncRNA of unknown function | Similar to *S. aureus* RsaK | 113 |
| 1170746-1170923 (+)<sup>‡</sup> | SPV_2317 |  | *srf-19* | ncRNA of unknown function |  | 55 |
| **1528520-1528643 (-)** | SPV_2378 |  | *srf-21* | ncRNA of unknown function |  |  |
| 1548804-1549088 (-) | SPV_2383 |  | *srf-22* | ncRNA of unknown function |  | 55 |
| 1598326-1598522 (+) | SPV_2392 |  | *ssrS* | 6S RNA |  | 53, **54** |
| 1873736-1873781 (-) | SPV_2433 |  | *srf-24* | ncRNA of unknown function |  | 54 |
| 1892857-1893007 (-) | SPV_2436 |  | *srf-25* | ncRNA of unknown function |  | 53, **54** |
| **1949385-1949547 (+)** | SPV_2442 |  | *srf-26* | ncRNA of unknown function |  |  |
| 1973172-1973570 (-)<sup>‡</sup> | SPV_2447 |  | *srf-27* | Type I addiction module antitoxin, Fst family | Not detected in this study due to its size; annotation based on sequence similarity | 58 |
| 1973509-1973913 (-) | SPV_2449 |  | *srf-28* | Type I addiction module antitoxin, Fst family |  | 58 |
| **2008242-2008356 (-)** | SPV_2454 |  | *srf-29* | ncRNA of unknown function | Similar to *L. welshimeri* LhrC |  |
| **2020587-2020685 (-)** | SPV_2458 |  | *srf-30* | ncRNA of unknown function |  |  |

Until now, several small RNA features have been reliably validated by Northern blot in *S. pneumoniae* strains D39, R6 and TIGR4 (53–57). Excluding most validation reports by Mann et al. due to discrepancies found in their data, 34 validated sRNAs were conserved in D39V. Among the 63 features identified here, we recovered and refined the coordinates of 33 out of those 34 sRNAs, validating our sRNA detection approach.

One of the detected sRNAs is the highly abundant 6S RNA (Figure 4B, left), encoded by ssrS, which is involved in transcription regulation. Notably, both automated annotations (RAST and PGAP) failed to report this RNA feature. We observed two differently sized RNA species derived from this locus, probably corresponding to a native and a processed transcript. Interestingly, we also observed a transcript containing both *ssrS* and the downstream tRNA gene. The absence of a TSS between the two genes, suggests that the tRNA is processed from this long transcript (Figure 4B, right).

Other detected small transcripts include three type I toxin-antitoxin systems as previously predicted based on orthology (58). Unfortunately, previous annotations omit these systems. Type I toxin-antitoxin systems consist of a toxin peptide (SPV_2132/SPV_2448/SPV_2450) and an antitoxin sRNA (SPV_2131/SPV_2447/SPV_2449). Furthermore, SPV_2120 encodes a novel sRNA that is antisense to the 3’-end of *mutR1* (SPV_0144), which encodes a transcriptional regulator (Figure 4C) and might play a role in controlling the production of MutR1, a putative transcriptional regulator (59).

### The pneumococcal genome contains at least 29 RNA switches

Several small RNA fragments were located upstream of protein-encoding genes, without an additional TSS in between. This positioning suggests that the observed fragment may be a terminated RNA switch, rather than a functional RNA molecule. Indeed, when we compared expression profiles of 5’-UTRs (untranslated regions) and the gene directly downstream, across infection-relevant conditions (9), we found several long UTRs (>100 nt) with a significantly higher average abundance than their respective downstream genes (Figure 5A). Such an observation may suggest conditional termination at the end of the putative RNA switch. We queried RFAM and BSRD (24) databases with the 63 identified small RNA features, which allowed us to immediately annotate 27 RNA switches (**Supplementary Table S7**).

**Figure 5.**
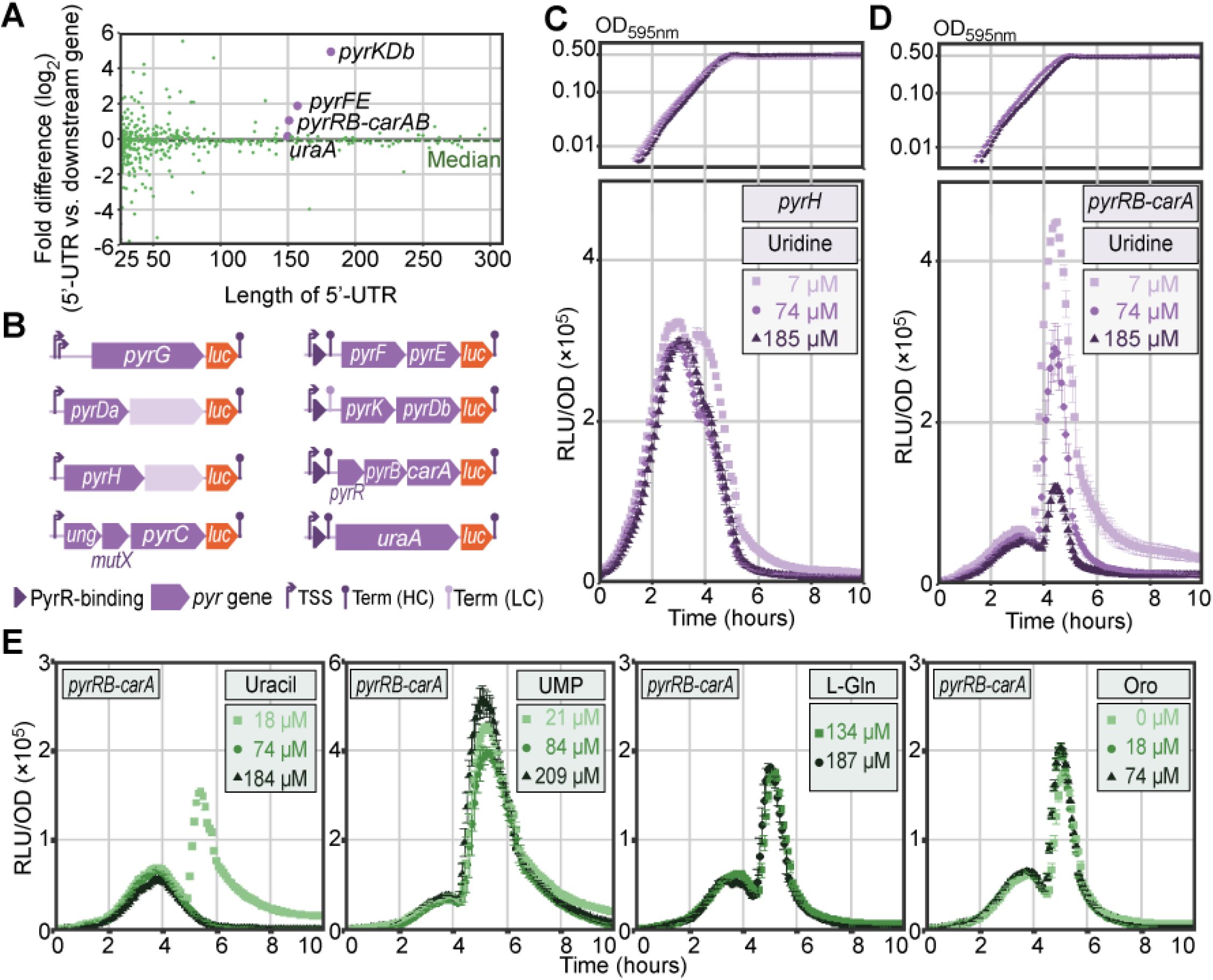
Long 5’-UTRs and validation of RNA switches. **A**. Mean ratio (^2^log) of 5’-UTR expression to that of the corresponding downstream gene. This fold difference was measured across 22 infection-relevant conditions (9) and plotted against 5’-UTR length. Among the overrepresented 5’-UTRs are several pyrimidine metabolism operons (*pyrKDb*, *pyrFE, pyrRB-carAB).* **B**. Firefly *luc* integration constructs used to assay the response to pyrimidine metabolites. Left: operons containing pyrimidine-related genes but lacking an upstream PyrR binding site. Right: pyrimidine operons with a predicted PyrR binding site (dark purple triangles). **C-D**. Growth (top) and normalized luciferase activity (RLU/OD, bottom) of *pyrH-luc* (C) and *pyrRB-carA-luc* (**D**) cells with varying uridine concentrations. **E**. Normalized luciferase activity (RLU/OD) of *pyrRB-carA-luc* cells with varying concentrations of (from left to right) uracil, uridine 5’-monophosphate, L-glutamine and orotic acid.

Furthermore, we found a candidate sRNA upstream of *pheS* (SPV_0504), encoding a phenylalanyl-tRNA synthetase component. Since Gram-positive bacteria typically regulate tRNA levels using so-called T-box leaders (60), we performed a sequence alignment between nine already identified T-box leaders and the *pheS* leader. Based on these results, we concluded that *pheS* is indeed regulated by a T-box leader (Figure 4D).

Finally, we identified a putative PyrR binding site upstream of *uraA* (SPV_1141), which encodes uracil permease. In a complex with uridine 5’-monophosphate (UMP), PyrR binds to 5’-UTR regions of pyrimidine synthesis operons, causing the regions to form hairpin structures and terminate transcription (61). This mechanism was already shown to regulate the expression of *uraA* in both Gram-positive (62) and Gram-negative (63) bacteria. Combining descriptions of PyrR in other species and sequence similarity with 3 other identified PyrR binding sites, we annotated this small RNA feature as a PyrR-regulated RNA switch, completing the 29 annotated RNA switches in D39V. To validate these annotations, we transcriptionally integrated a gene encoding firefly luciferase (*luc*) behind the four putative PyrR-regulated operons (*pyrFE*, *pyrKDb, pyrRB-carAB, uraA*), along with four negative controls (*pyrG*, *pyrDa-holA, pyrH, ung-mutX-pyrC*) that do contain genes involved in pyrimidine metabolism but lack a putative PyrR-binding site (Figure 5B). As expected, expression of operons lacking a PyrR RNA switch is not affected by increasing concentrations of uridine (for example in *pyrH-luc*, Figure 5C), while the putative PyrR-regulated operons are strongly repressed (for example in *pyrRB-carA-luc*, Figure 5D). Finally, we tested other intermediates from uridine metabolism and observed a similar trend when uracil was added instead. UMP exhibited only a marginal effect on the expression of the *pyrR* operon, with much weaker repression than observed with comparable uridine concentrations. This may be explained by the fact that UMP itself is not efficiently imported by most bacteria (64, 65), but first needs to be dephosphorylated extracellularly. Potential absence or inefficiency of the latter process in *S. pneumoniae* could explain this observation. Other intermediates in the pyrimidine metabolism, L-glutamine and orotic acid, did not incite an observable effect on *pyrR* expression. Interestingly, the PyrR-binding regulatory element was recently shown to be essential for survival, successful colonization and infection in mice (BioRxiv: https://doi.org/10.1101/286344).

### Combining putative termination peaks and predicted stem-loops to annotate terminators

Aside from the customary annotation of gene and pseudogene features, we complemented the annotation with transcriptional regulatory elements including TSSs, terminators and other transcription regulatory elements. First, we set out to identify transcriptional terminators. Since *S. pneumoniae* lacks the Rho factor (66), all of its terminators will be Rho-independent. We combined the previously generated list of putative termination peaks with a highly sensitive terminator stem-loop prediction by TransTermHP (22) and annotated a terminator when a termination peak was found within 10 bps of a predicted stem loop. The genome-wide distribution of the 748 annotated terminators is shown in Figure 6A. Terminator characteristics (**Supplementary Figure S6**) resembled those observed in *B. subtilis* and *E. coli* (67). We further compared the number of sequenced fragments (i.e. molecules in the RNA-seq library) ending at each termination peak with the number of fragments covering the peak without being terminated. This allowed us to estimate the termination efficiency of each terminator in the genome, which is listed for all terminators in **Supplementary Table S8** and visible in PneumoBrowse (see below).

### Direct enrichment of primary transcripts allows precise identification of transcription start sites

We determined TSSs using a novel technique (Cappable-seq, (7)): primary transcripts (i.e. not processed or degraded) were directly enriched by exploiting the 5’-triphosphate group specific for primary transcripts, in contrast with the 5’-monophosphate group that processed transcripts carry. We then sequenced the 5’-enriched library, along with a nonenriched control library, and identified 981 TSSs with at least 2.5-fold higher abundance in the enriched library than in the untreated control library. Taking into account the possibility of rapid 5’-triphosphate processing, we added 34 (lower confidence, LC) TSSs that were not sufficiently enriched, yet met a set of strict criteria (see **Materials and Methods** for details). The genomic distribution of these 1,015 TSSs is shown in Figure 6A. Importantly, our data could closely reproduce previously reported TSSs (**Supplementary Table S9**), as determined by 5’-RACE or primer extension experiments (38, 56, 68–86). Out of 54 experimentally determined TSSs, 43 were perfectly reproduced by our data, 6 were detected with an offset of 1 or 2 nucleotides, and 4 were visible in the raw data tracks, but did not meet our detection criteria. Finally, for operon *groESL*, Kim et al. reported a TSS 155 nucleotides upstream of the *groES* TIS (87), while our data contained zero reads starting at that site. Instead, we detected a very clear TSS 62 nucleotides upstream of *groES*, preceded by an RpoD-binding site and closely followed by a CIRCE element (88), which serves as a binding site for heat-shock regulator HrcA. We found that this promoter architecture matches that observed in *Lactococcus lactis* (S.B. van der Meulen and O.P. Kuipers, unpublished), *Geobacillus thermoglucosidasius* (E. Frenzel, O.P. Kuipers and R. van Kranenburg, unpublished) and *Clostridium acetobutylicum* (89), validating the TSS reported here. These results confirm that the we used conservative detection criteria and that we only report TSSs with a high degree of confidence.

### Strong nucleotide bias on transcriptional start sites

Analysis of the nucleotide distribution across all TSSs showed a strong preference for adenine (A, 63% vs. 30% genome-wide) and, to a lesser extent, guanine (G, 28% vs. 22%) on +1 positions (Figure 6B). While it has to be noted that especially upstream regions are biased by the presence of transcriptional regulatory elements (see below), we observed a general bias around the TSSs towards sequences rich in adenine and poor in cytosine (C) (Figure 6C). A striking exception to this rule is observed on the −1 position, where thymine (T) is the most frequently occurring base (51%) while A is underrepresented (16%). Interestingly, while *Escherichia coli* (7) and *Salmonella enterica* (90) qualitatively show the same preference for adenines on the +1 position and against adenines on the −1 position, these biases seem to be absent in *Helicobacter pylori* (91).

**Figure 6.**
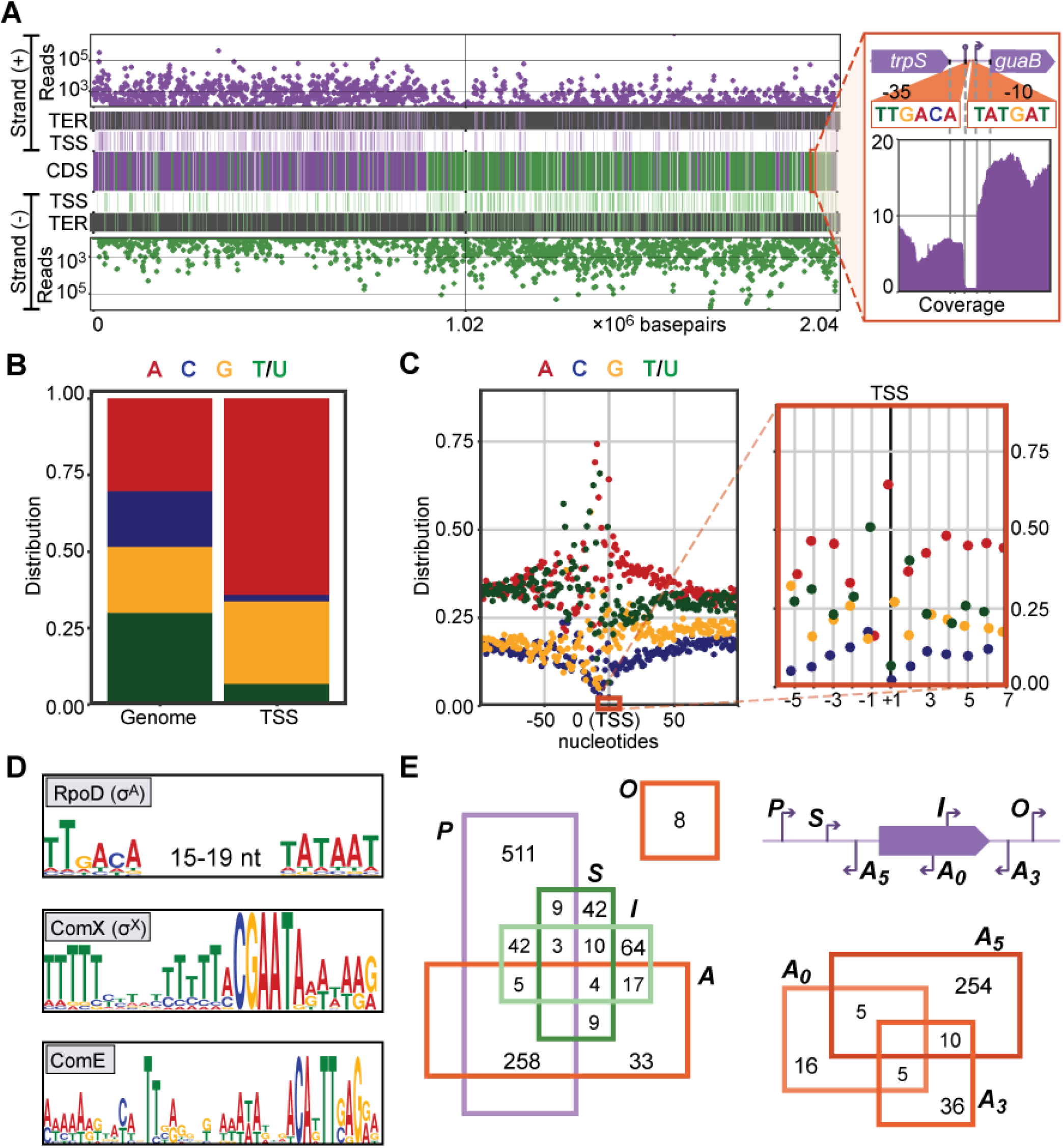
Characterization of transcriptional start sites. **A**. Genome-wide distributions of sequencing reads, terminators (TER) and transcriptional start sites (TSS) on the positive (top) and negative strand (bottom) and annotated coding sequences (middle) are closely correlated. Features on the positive and negative strand are shown in purple and green, respectively. The inset shows the coverage in the *trpS-guaB* locus, along with a detected terminator (84% efficient), TSS, and RpoD-binding elements (−35 and −10). **B**. Nucleotide utilization on +1 (TSS) positions, compared to genome content. **C**. Nucleotide utilization around TSSs (left: −100 to +101, right: −6 to +7). **D**. RpoD (σ_A_), ComX (σ_X_) and ComE binding sites found upstream of TSSs. E. TSSs were divided, based on local genomic context, into five classes (top right): primary (P, only or strongest TSS within 300 nt upstream of a feature), secondary (S, within 300 nt upstream of a feature, not the strongest), internal (I, inside a feature), antisense (A), and orphan (O, in none of the other classes). Results are shown in a Venn diagram (left). Antisense TSSs were further divided into 3 subclasses (bottom right): A_5_ (upstream of feature), A_3_ (downstream of feature), and A_0_ (inside feature).

### Promoter analysis reveals regulatory motifs for the majority of transcription start sites

We defined the 100 bps upstream of each TSS as the promoter region and used the MEME Suite (92) to scan each promoter region for regulatory motifs (Figure 6D: see **Supplementary Methods** for detailed constraints). We could thereby predict 382 RpoD sites (σ_A_, (93)), 19 ComX sites (σ_X_, (94)) and 13 ComE sites (95). In addition to the complete RpoD sites, another 449 TSSs only had an upstream −10 (93) or extended −10 sequence (96). Finally, we annotated other transcription-factor binding sites, including those of CodY and CcpA, as predicted by RegPrecise (97).

### Characterization of TSSs based on genomic context and putative operon definition

Subsequently, the TSSs were classified based on their position relative to annotated genomic features (91), categorizing them as primary (*P*, the only or strongest TSS upstream of feature), secondary (*S*, upstream of a feature, but not the strongest TSS), internal (*I*, inside annotated feature), antisense (*A*, antisense to an annotated feature) and/or orphan (O, not in any of the other categories), as shown in Figure 6E. Notably, we categorized antisense TSSs into three classes depending on which part of the feature the TSS overlaps: *A_5_* (in the 100 bps upstream), *A_0_* (within the feature), or *A_3_* (in the 100 bps downstream). Doing so, we could classify 828 TSSs (81%) as primary (pTSS), underscoring the quality of TSS calls. Finally, we defined putative operons, starting from each pTSS. The end of an operon was marked by (i) the presence of an efficient (>80%) terminator, (ii) a strand swap between features, or (iii) weak correlation of expression (<0.75) across 22 infection-relevant conditions (9) between consecutive features. The resulting 828 operons cover 1,391 (65%) annotated features (**Supplementary Table S10**) and are visualized in PneumoBrowse.

### Coding sequence leader analysis reveals ribosome-binding sites and many leaderless genes

We evaluated the relative distance between primary TSSs and the first gene of their corresponding operons (Figure 7A). Interestingly, of the 768 operons starting with a coding sequence (or pseudogene), 81 have a 5’-UTR too short to harbor a potential Shine-Dalgarno (SD) ribosome-binding motif (leaderless; Figure 7A, inset). In fact, for 70 of those (9% of all operons), transcription starts exactly on the first base of the coding sequence, which is comparable to findings in other organisms (98, 99). Although the presence of leaderless operons suggests the dispensability of ribosomal binding sites (RBSs), motif enrichment analysis (92) showed that 69% of all coding sequences do have an RBS upstream. To evaluate the translation initiation efficiency of leaderless coding sequences, we selected the promoter of leaderless *pbp2A* (SPV_1821) and cloned it upstream of a reporter cassette, containing *luc* and *gfp.* The cloning approach was threefold: while *gfp* always contained an upstream RBS, *luc* contained either (i) no leader, (ii) an upstream RBS, or (iii) no leader and a premature stop codon (Figure 7B). Fluorescence signals were comparable in all three constructs, indicating that transcription and translation of the downstream *gfp* was unaffected by the different *luc* versions (Figure 7C). Bioluminescence assays (Figure 7D) showed, surprisingly, that the introduction of an upstream RBS had no effect on luciferase production. Although it is unknown why translation efficiency of this leaderless gene is completely insensitive to the introduced RBS, one could speculate that transcription and translation are so efficiently coupled, that an RBS becomes redundant. Additionally, the absence of signal in the strain with an early stop codon allowed us to rule out potential downstream translation initiation sites. This experiment provides direct evidence that leaderless genes can be efficiently translated.

**Figure 7.**
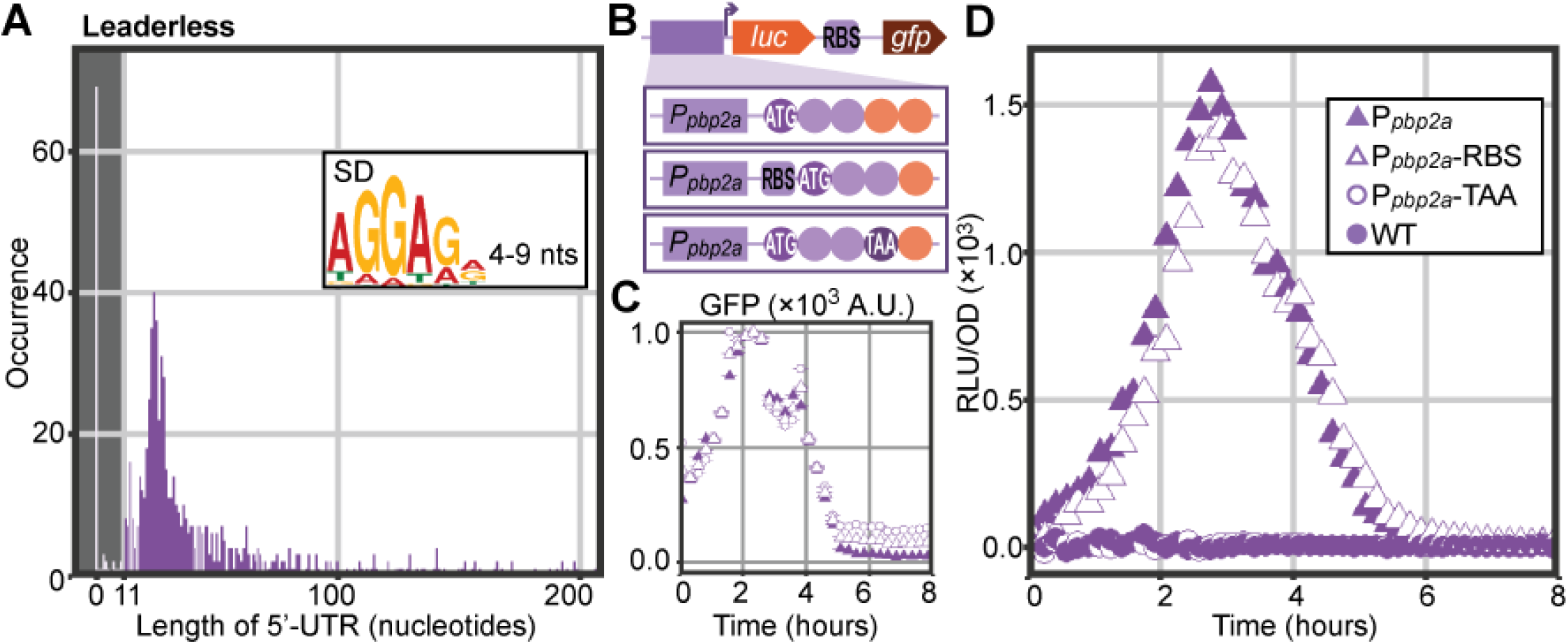
Leaderless coding-sequences and pseudogenes. **A**. Length distribution of 5’-untranslated regions of all 768 operons that start with a CDS or pseudogene shows 70 (9%) leaderless transcripts. The shaded area contains leaders too short to contain a potential Shine-Dalgarno (SD) motif (inset). **B**. In an ectopic locus, we cloned the promoter of leaderless gene *pbp2a* upstream of an operon containing *luc* (firefly luciferase) and *gfp* (superfolder GFP), the latter always led by a SD-motif (RBS). Three constructs were designed: (i) native promoter including the first three codons of *pbp2a*, (ii) the same as (i), but with an additional RBS in front of the coding sequence, (iii) the same as (i), but with a stop codon (TAA) after three codons. **C**. Fluorescence signal of all three constructs shows successful transcription. **D**. Normalized luminescence signals in all three strains and a wild-type control strain. Successful translation of *luc* is clear from luminescence in P_*pbp2a*_-*luc* (filled triangles) and adding an RBS had no measurable effect on luciferase activity (open triangles). Integration of an early stop codon (open circles) reduced signal to wild-type level (filled circles).

### PneumoBrowse integrates all elements of the deep D39V genome annotation

Finally, we combined all elements of the final annotation to compile a user-friendly, uncluttered genome browser, PneumoBrowse (https://veeninglab.com/pneumobrowse, Figure 8). Based on JBrowse (8), the browser provides an overview of all genetic features, along with transcription regulatory elements and sequencing coverage data. Furthermore, it allows users to flexibly search for their gene of interest using either its gene name or locus tag in D39V, D39W or R6 (prefixes ‘SPV_’, ‘SPD_’ and ‘spr’, respectively). Right-clicking on features reveals further information, such as its gene expression profile across 22 infection-relevant conditions and co-expressed genes (9).

**Figure 8.**
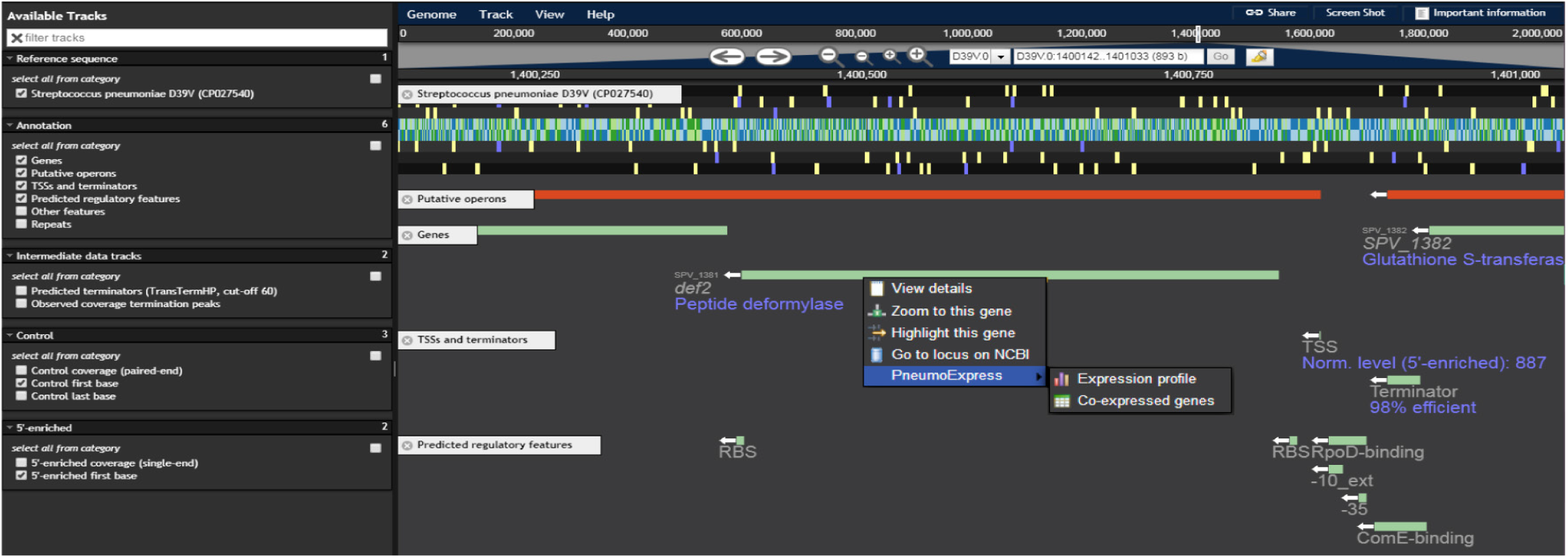
PneumoBrowse. A screenshot of the *def2* locus in PneumoBrowse shows the richness of available data. In the left pane, tracks can be selected. In the right pane, annotation tracks (*Putative operons, Genes, TSSs and terminators, Predicted regulatory features*) are shown along with data tracks (*Control first base, 5-enriched first base).* Regulatory features annotated upstream of *def2* include a 98% efficient terminator, ComE- and RpoD-binding sites, a TSS and an RBS. Additionally, a context menu (upon right-click) provides links to external resources, such as PneumoExpress (9).

## DISCUSSION

Annotation databases such as SubtiWiki (100) and EcoCyc (101) have tremendously accelerated gene discovery, functional analysis and hypothesis-driven research in the fields of model organisms *Bacillus subtilis* and *Escherichia coli*. It was therefore surprising that such a resource was not yet available for the important opportunistic human pathogen *Streptococcus pneumoniae*, annually responsible for more than a million deaths (102). With increases in vaccine escape strains and antimicrobial resistance, a better understanding of this organism is required, enabling the identification of new vaccine and antibiotic targets. By exploiting recent technological and scientific advances, we now mapped and deeply annotated the pneumococcal genome at an unprecedented level of detail. We combined state-of-the-art technology (e.g. SMRT sequencing, Cappable-seq, a novel sRNA detection method and several bioinformatic annotation tools), with thorough manual curation to create PneumoBrowse, a resource that allows users to browse through the newly assembled pneumococcal genome and inspect encoded features along with regulatory elements, repeat regions and other useful properties (Figure 8). Additionally, the browser provides direct linking to expression and co-expression data in PneumoExpress (9). The methods and approaches applied here might serve as a roadmap for genomics studies in other bacteria. It should be noted that the detection of sRNAs and transcription regulatory elements, such as TSSs and terminators, relies on active transcription of these features in the studied conditions. Therefore, new elements may still be elucidated in the future. The D39V annotation will periodically be updated with information from complementary studies to ours, such as recent work by Warrier et al. (BioRxiv: https://doi.org/10.1101/286344), once peer-reviewed.

We showed that, like D39W, D39V strongly resembles the ancestral strain NCTC 7466, although it has lost the cryptic plasmid pDP1 and contains a number of single-nucleotide mutations. Several differences were observed with the sequence of D39W, a separate isolate of NCTC 7466 (**Supplementary Figure S1**), as reported 11 years ago ((6), Table 1). Upon comparison to several other pneumococcal strains, we identified multiple large chromosomal inversions (Figure 2). Although the inversion observed in D39V contained the terminus of replication, it created an asymmetry in replichore length, visible both in the location of the dif/Xer locus (Figure 3A) and the direction of encoded features (Figure 6A). Such inversions are likely to be facilitated by bordering repeat regions. In this context, the many identified repeat regions (i.a. BOX elements, RUPs and IS elements) may play an important role in pneumococcal evolution, by providing a template for intragenomic recombination or inversion events (103, 104). For example, Rued et al. identified four strains that suppressed the lethal effect of the deletion of *gpsB* (SPV_0339), encoding a regulatory protein that is likely involved in the coordination of septal and peripheral cell wall synthesis (105). Two of these suppressors contained a large deletion bordered by an even larger duplication. Intriguingly, these large genetic changes occurred in the exact genomic region that was found to be inverted in D39V. Although the exact mechanism behind these mutations is unclear, they are likely to be facilitated by the inverted repeats surrounding the affected region.

We further unveiled the mosaic nature of pneumococcal histidine triad proteins, and especially vaccine candidate PhtD, variation of which is facilitated both by the aforementioned chromosomal inversion and more local recombination events. This observation expands the set of previously identified highly variable pneumococcal antigens (106, 107).

Despite the differences observed with NCTC 7466, D39V is genetically stable under laboratory conditions, as both the *hsdS* locus and the *ter* configuration are identical in all analyzed derivative strains (*not shown).* Additionally and importantly, D39V is virulent in multiple infection models (108–111), making it an ideal, stable genetic workhorse for pneumococcal research, along with D39W and NCTC 7466. Therefore, strain D39V was made available to the community through the Public Health England (PHE) National Collection of Type Cultures (NCTC 14078).

Among the annotated 1,877 CDSs, 12 rRNAs, 58 tRNAs and 165 pseudogenes, there are 89 CDSs and 109 pseudogenes previously not annotated (**Supplementary Tables S4-S6**). Additionally, we identified and annotated 34 sRNAs and 29 RNA switches (Table 2, Supplementary Table S7, Figure 5). We look forward to seeing future studies into the function of the newly identified proteins and sRNAs. The novel sRNA detection method employed, based on fragment size distribution in paired-end sequencing data (Figure 4), while already sensitive, can probably be further improved by employing sRNA-specific library techniques, combined with more sophisticated statistical analyses.

Finally, to understand bacterial decision-making, it is important to obtain detailed information about gene expression and regulation thereof. Therefore, we identified transcriptional start sites and terminators (Figure 6), sigma factor binding sites and other transcription-regulatory elements. This architectural information, complemented by expression data from PneumoExpress (9), will be invaluable to future microbiological research and can be readily accessed via PneumoBrowse.

## AVAILABILITY

Python scripts used for data analysis are available at https://github.com/veeninglab/Spneu-deep-annotation. PneumoBrowse can be accessed via https://veeninglab.com/pneumobrowse. All detected and annotated small RNA features were submitted to the Bacterial Small Regulatory RNA Database (BSRD).

## ACCESSION NUMBERS

SMRT sequencing data used for de novo assembly was deposited to SRA (SRP063763). Unless stated otherwise, RNA-seq datasets used in this study can be found in the SRA repository (SRP133365). The D39V assembly and annotation are accessible at GenBank (CP027540).

## ACKNOWLEDGEMENTS

We are grateful to A. Patrignani (Functional Genomics Center Zurich) for support in SMRT sequencing; F. Thümmler (vertis Biotechnologie AG) for support in Illumina RNA-seq; A. de Jong and S. Holsappel for (bio)informatics support; J. Donner for kindly providing DNA-seq data for the three investigated clinical isolates; and S.B. van der Meulen and E. Frenzel for sharing their TSS data.

