## Supplementary Materials for "Deep genome annotation of the opportunistic human pathogen *Streptococcus pneumoniae* D39" (from another version of this preprint: version 2, 2018-03-22)

**SUPPLEMENTARY FIGURES**

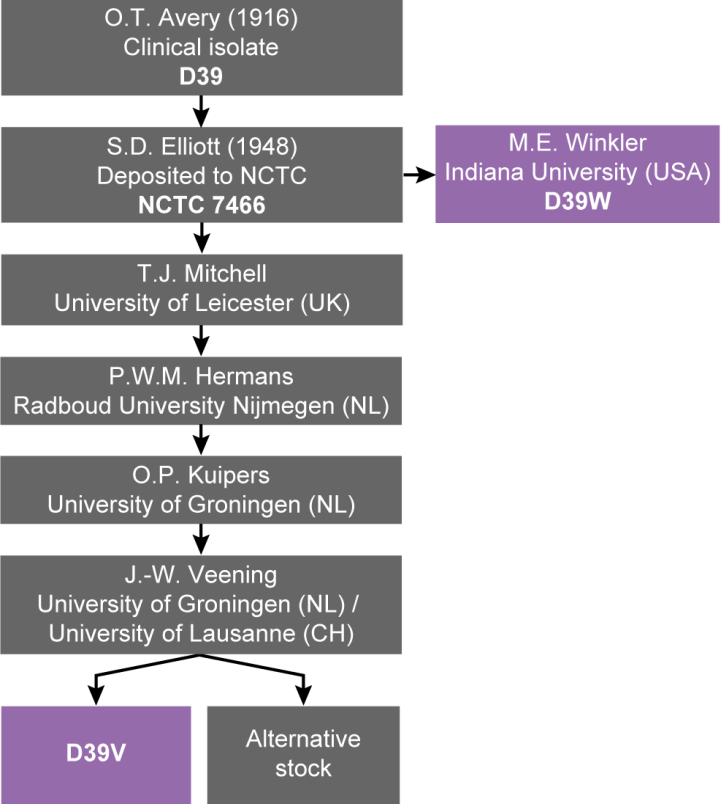

**Figure S1.** History of D39V. D39 was isolated in 1916 by Avery and submitted to the National Collection of Type Cultures (NCTC) in 1948. From there, cells have been recultured several times before reaching the Veening lab (D39V).

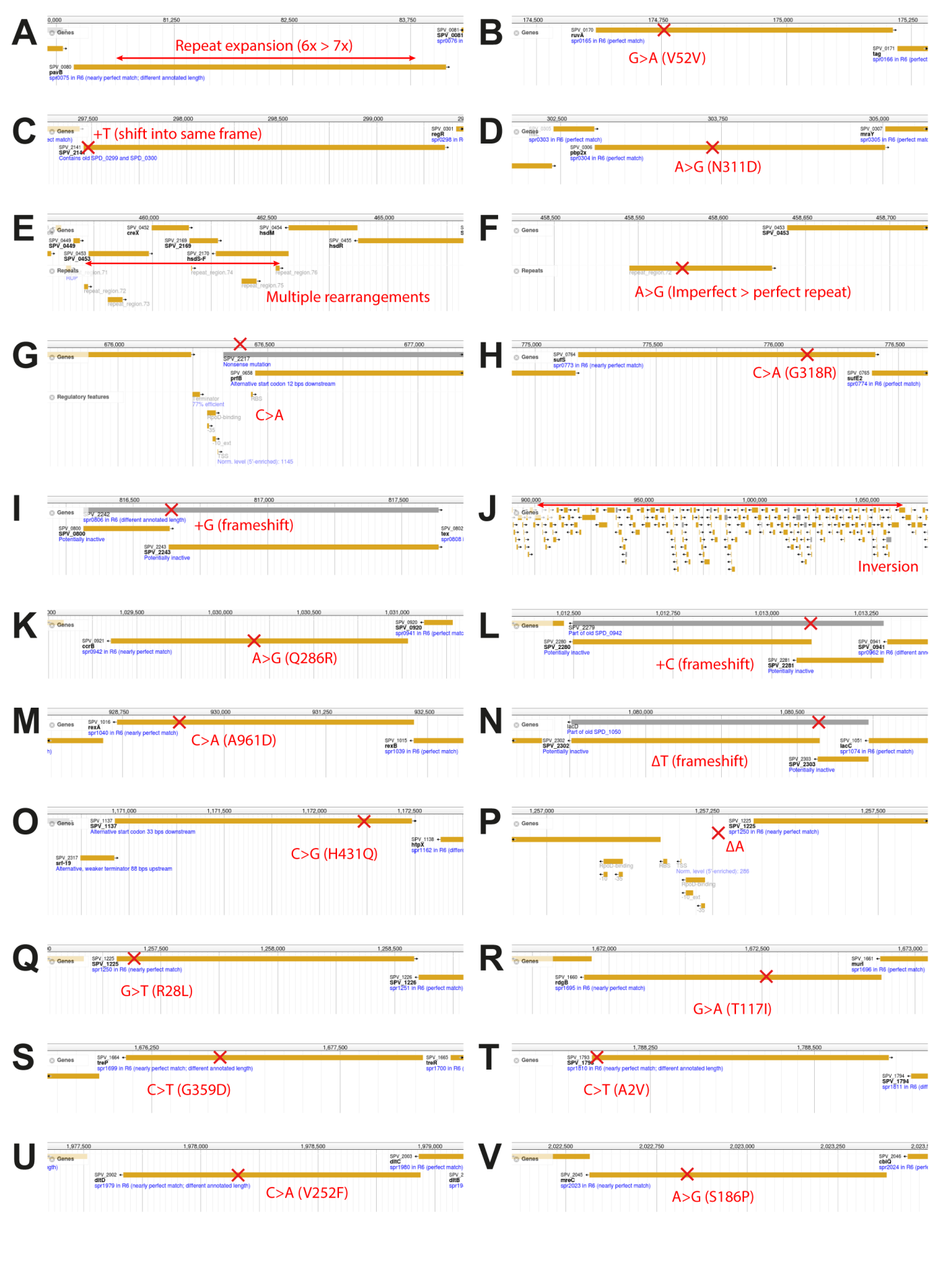
**Figure S2.** Genomic context in D39V of sites of differing sequence with D39W (CP000410). A-V. Regions containing sequence differences, matching the order in main Table 1. The exact location of observed differences and their consequences, based on the situation in the old assembly, are indicated in red.

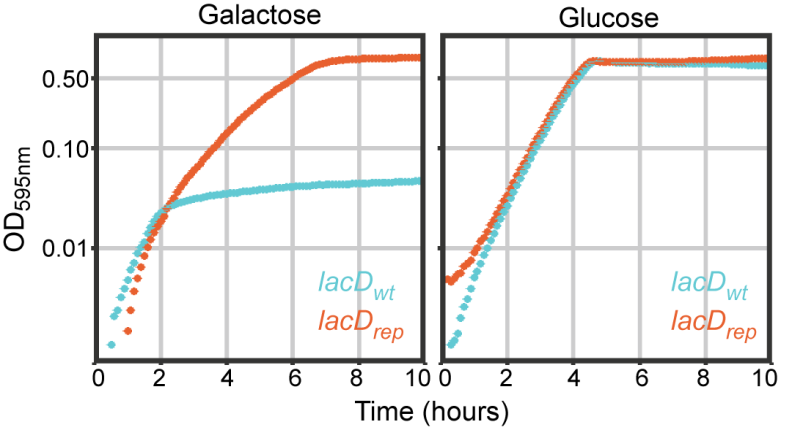

**Figure S3.** Restored growth on galactose of a *lacD*-repaired strain. Wild-type D39V (*wt*) and a mutant carrying the intact R6 allele of *lacD* (*rep*), were grown in a plate reader in C+Y medium with either galactose (left) or glucose (right) as primary carbon source. As expected, repairing *lacD* restored growth on galactose.

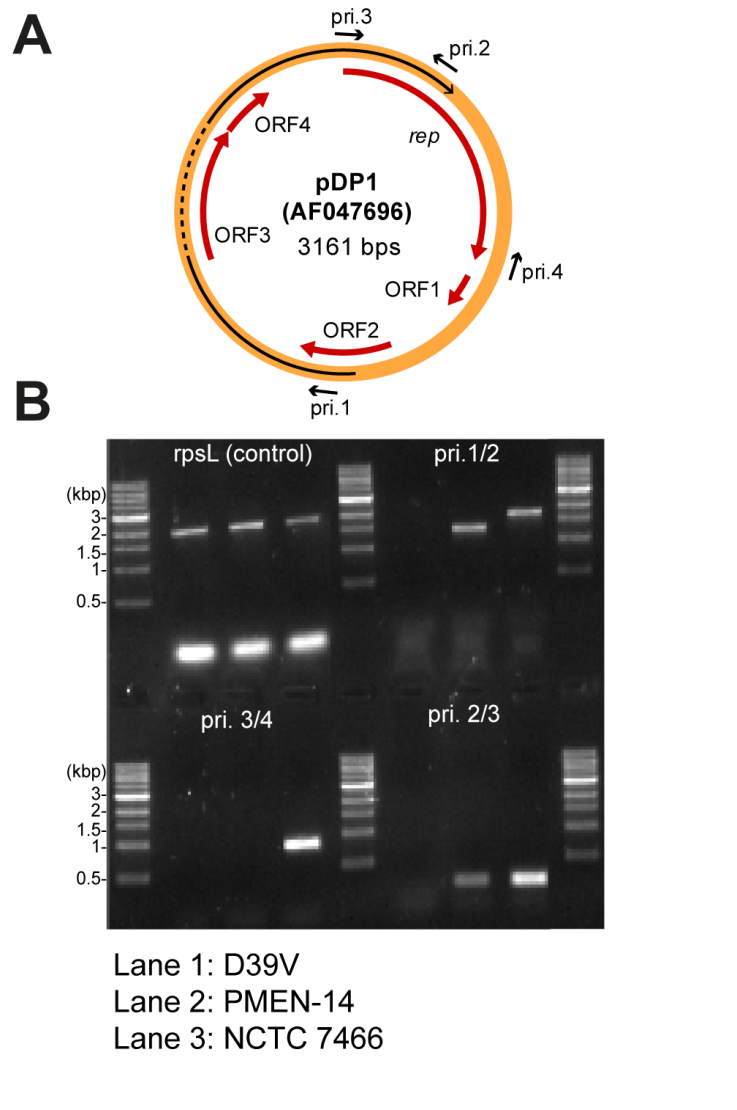

**Figure S4.** PCR-based assay on presence or absence of cryptic plasmid pDP1. A. Organization of plasmid pDP1 (orange), including binding sites of primers used for amplification. The black line indicates how part of the plasmid was integrated in strain PMEN-14. B. Electrophoresis gel after DNA amplification with 4 primer pairs. Lanes 1-3: D39V, PMEN-14 and NCTC 7466. Positive control (*rpsL*) gives the correct product for all samples. All expected products are observed for both NCTC 7466 samples and for PMEN-14, which has a degenerate version of the plasmid integrated into its chromosome. The absence of product in all of the D39V assays confirms that this strain lacks plasmid pDP1.

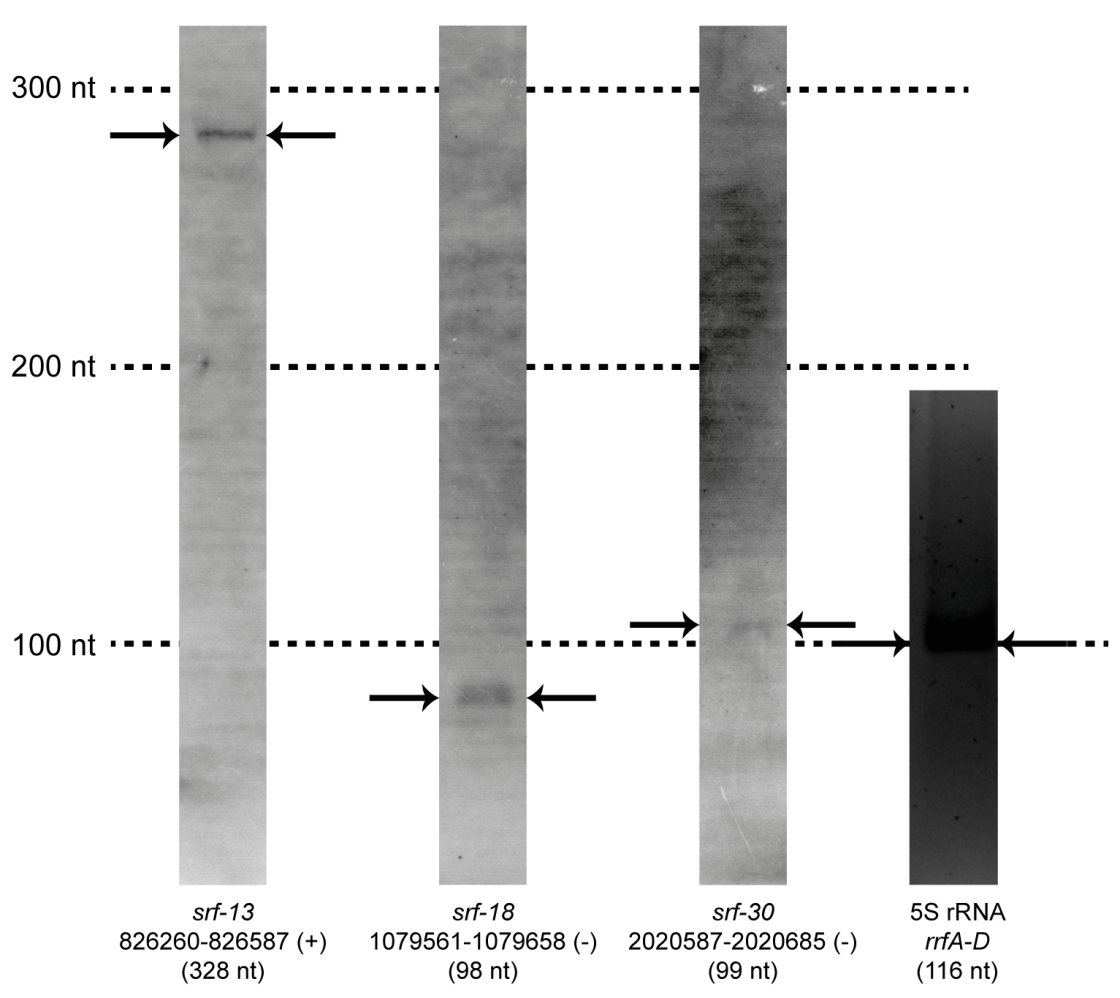

**Figure S5.** Northern blot analysis. The presence and size of three identified small RNA features (*srf-13*, *srf-18*, *srf-30*) was confirmed by Northern blot analysis. As a positive control, 5S ribosomal RNA was also included in the analysis (right).

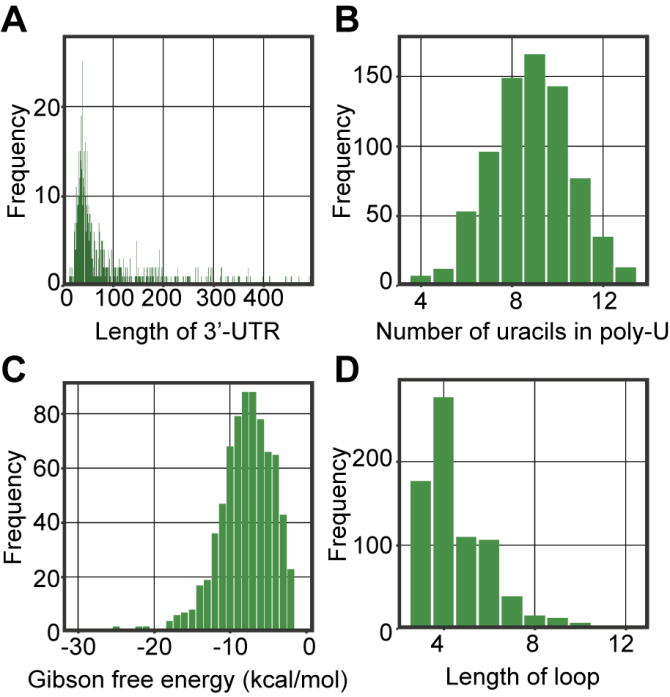

**Figure S6.** Characteristics of annotated terminators. Only terminators were included that both contained a predicted stem-loop (TransTermHP, (1)) and a detected termination peak. A. Distribution of terminator distance to the first feature upstream (3’-UTR). B. Frequency distribution of the number of uracils in a 15-nucleotide window following the predicted stem-loop structure. C. Binding energy distribution of the stem sequence, as reported by TransTermHP. D. Length distribution of the loop between both arms of the terminator stem.

**SUPPLEMENTARY METHODS**

**Culturing of *Streptococcus pneumoniae* D39 and transformation**

*S. pneumoniae* strains were routinely grown at 37°C, diluted 1:100 from -80°C stock into C+Y (pH 6.8), a liquid casein-based medium, supplemented with yeast extract (Sigma-Aldrich, Boom, NL), as described in the accompanying paper by Aprianto et al. (BioRxiv: <https://doi.org/10.1101/283739>).

Construction of strains was performed as previously described (2). For transformation, cells were grown until OD_600nm_ ~0.1. Subsequently, 100 ng·ml^-1^ synthetic CSP-1 (competence-stimulating peptide-1) was added into the suspension and incubation was continued for another 12 min at 37°C. Transforming DNA was added to the now-competent *S. pneumoniae*. Cells were then incubated at 30°C for 20 min to allow for DNA internalization. Afterwards, fresh C+Y was added and the suspension was incubated further at 37°C for 90 minutes. Selection of transformants was done by plating in Columbia agar (Oxoid, UK) supplemented with 2% sheep blood (Johnny Rottier, NL) and 0.50 μg·ml^-1^ tetracycline for *luc-gfp* constructs or 0.25 mg·ml^-1^ kanamycin for *luc* constructs. The following day, colonies were streaked

on antibiotic-supplemented blood agar and incubated overnight in 5% (v/v) CO_2_. Constructs were checked by colony PCR and Sanger sequencing.

**Strain construction**

*Transcriptional fusions of pyrimidine operons and* luc. Strains (RA59, RA60, RA61, RA62, RA63, RA65, RA68, RA69) were constructed in which *luc* was inserted downstream of each of eight different pyrimidine metabolism operons (*pyrG*, *pyrFE*, *pyrDa*-*holA*, *pyrH*-*frr*, *pyrKDb*, *ung*-*mutX*-*pyrG*, *pyrRB*-*carA*, *uraA*). For each operon, upstream and downstream fragments were PCR-amplified from genomic DNA, using primer pairs [last_gene]_up_F / [last_gene]_up_R_BglII and [last_gene]_down_F_NotI / [last_gene]_down_R, respectively. The cassette carrying *luc* and a kanamycin resistance marker was amplified from strain MK134 with primer pair Luc_F_BglII_AseI_EcoRI / kan-R+NotI. Fragments were cut with BglII and NotI and ligated together. Subsequently, wild-type D39V cells were transformed with the ligation product and positive clones were selected on kanamycin and verified by Sanger sequencing.

*Transcriptional fusions of* *P*_pbp2a_ *and* luc.	Three strains were constructed in which *luc* was controlled by P*_pbp2a_* in an ectopic locus (*bgaA*). In RA114, the leader of *luc* contained just the first three codons of *pbp2a*. Plasmid pLA18 (*luc*_*gfp*) was amplified with primer pair G033 / G044, while P*_pbp2a_* was amplified with G035 / G036. In RA115, the *luc* leader also contained an RBS. Plasmid pLA18 was amplified with G037 / G034, while P*_pbp2a_* was amplified with G035 / G038. In RA122, the *luc* leader contained the first three codons of *pbp2a* followed by a stop codon. Plasmid pLA18 was amplified with G039 / G034, while P*_pbp2a_* was amplified with G035 / G040. For each strain, both fragments were assembled using Gibson’s assembly (3) and the resulting molecule was used to transform wild-type D39V cells. Positive clones were selected on tetracycline and verified by Sanger sequencing.

*Strain with repaired* lacD. To repair the observed frameshift in *lacD*, we replaced the D39V allele of *lacD* with the intact version of the gene from *S. pneumoniae* R6. The latter gene was amplified from R6 cells with primer pair lacD_F / lacD_R and the PCR product was used to transform D39V cells. Positive clones were identified by growth on galactose and further verified by Sanger sequencing.

**Growth, luciferase and GFP assays**

Before the assays, cells were routinely pre-cultured in C+Y medium until an OD_600_ of 0.4, and then diluted 1:100 into fresh medium in a 96-wells plate. All assays were performed in a Tecan Infinite 200 PRO at 37°C.

Luciferase assays were performed in C+Y with 0.25 mg·ml^-1^ D-luciferin, sodium salt and signals were normalized through division by OD_595_.

To normalize fluorescence signals, we included a parental strain without *gfp* and used the growth-dependent autofluorescence readings of that strain to subtract from fluorescence signals of *gfp*-containing strains.

Growth assays of *lacD*-repaired strains were performed in C+Y with either 10.1 mM galactose or 10.1 mM glucose as main carbon source.

**DNA and RNA isolation, primary transcript enrichment and sequencing**

Wild-type *S. pneumoniae* cells were pre-cultured in C+Y medium and diluted, 1:100, into 100 ml fresh C+Y medium. Cells were harvested by centrifugation at an OD_600_ of 0.4. Isolation of genomic DNA was performed using the Wizard Genomic DNA Purification Kit (Promega), where the nuclei lysis buffer was supplemented with 0.05% SDS, 0.025% sodium deoxycholate and 200 mg/ml RNase A. DNA was further purified by ethanol precipitation. A total of 12 μg genomic DNA was sent to the Functional Genomics Center Zurich, where a 6/8 kbp insert library was created with a lower cut-off of 4kbp and was then sequenced using a PacBio RS II machine. Supporting Illumina HiSeq DNA-seq data was obtained previously as described (4, 5).

D39V samples for RNA-seq were pre-cultured in suitable medium before inoculation (1:100) into four infection-relevant conditions: lung-like medium, CSF-like medium, fever-like conditions in CSF-like medium and late competence (20 min after CSP addition) in C+Y medium. Composition of media, a detailed description of conditions and the total RNA isolation protocol are described in the accompanying paper by Aprianto et al. (BioRxiv: <https://doi.org/10.1101/283739>). Isolated RNA was sent to vertis Biotechnologie AG. Total RNA from the four conditions was combined in an equimolar fashion and the pooled RNA was divided into two portions. The first portion was directly enriched for primary transcripts (Cappable-seq (6)) and, after library preparation according to the manufacturer’s recommendations, sequenced on Illumina NextSeq in single-end (SE) mode. This technique exploits the fact that primary transcripts carry a 5’-triphosphate group, while processed transcripts carry a monophosphate group. The second RNA portion was rRNA-depleted and sequenced on Illumina NextSeq in paired-end (PE) mode, to obtain information about both 5’ and 3’ ends of sequenced fragments.

***De novo* assembly of the D39V genome and DNA modification analysis**

*De novo* genome assembly was performed using the Hierarchical Genome Assembly Process (HGAP3) module of the PacBio SMRT portal version 2.3.0. This resulted in two contigs: one of over 2 Mbp with 250-500x coverage, and one of 12 kbp with 5-25x coverage. The latter, small contig was discarded based on its low coverage and high sequence similarity with a highly repetitive segment of the larger contig. The large contig was circularized manually, by removing a 14 kbp sequence that occurred both at the start and end of the sequence. The resulting sequence was verified by re-mapping the data using the ‘RS_Resequencing.1’ module and then rotated such that *dnaA* was positioned on the positive strand, starting on the first nucleotide. Previously published Illumina data (GEO accessions GSE54199 and GSE69729) were mapped on the new assembly, using *breseq* (7). The datasets lacked coverage in the *bgaA* locus, because the sequenced strains were *bgaA*::P*_ssbB_*-*luc*. This locus, together with four loci of potential mistakes in the assembly were verified by Sanger sequencing. Only one actual mistake was identified in the *de novo* assembly.

DNA modification analysis was performed using the ‘RS_Modification_and_Motif_Analysis.1’ module in the SMRT portal, with a QV cut-off of 100.

**Detection of cryptic plasmid pDP1**

To evaluate whether an analogous plasmid to pDP1 was present in D39V, we performed a PCR-based assay. Amplification products were obtained for NCTC 7466 cells, with primer pairs pDP1.1F / pDP1.2R; pDP1.3F / pDP1.4R; and pDP1.3F / pDP1.2R. The absence of these products when using D39V as a template suggested that no such plasmid existed in this strain. As an extra validation step, we mapped RNA-seq data to the DNA sequence of pDP1, using Bowtie 2 (8). When no reads mapped to pDP1, we concluded that the plasmid was indeed absent from D39V.

**Automated and curated annotation**

The assembled genome sequence was annotated automatically, using PGAP ((9), executed October) 2015) and RAST ((10), executed June 2016). The results of both annotations were compared and for each discrepancy, support was searched. Among the support used were scientific publications (PubMed), highly similar features (BLAST (11)), reviewed UniProtKB entries (12) and/or detected conserved domains as found by CD-Search (13). When no support was found for either the PGAP or RAST annotation, the latter was used. Duplicate gene names were also resolved during curation. After the first round of curation, the resulting annotation was compared with the curated annotation of R6 (NC_003098.1) to complete the working annotation by identifying loci potentially mis-annotated by RAST and PGAP. R6 is a D39 derivative that shares many identical features. Again, the R6 annotation was only adopted in the presence of convincing support. Finally, an extensive literature search was performed, with locus tags and (if available) gene names from the old D39 annotation (prefix: ‘SPD_’) as query. When identical features were present in R6, a similar search was performed with R6 locus tags (prefix ‘spr’) and gene names. Using the resulting literature, the annotation was further refined.

CDS pseudogenes were detected by performing a BLASTX search against the NCBI non-redundant protein database, using the DNA sequence of two neighboring genes and their intergenic region as query. If the full-length protein was found, the two (or more, after another BLASTX iteration) genes were merged into one pseudogene.

Furthermore, sRNAs and riboswitches, transcriptional start sites and terminators, transcription-regulatory sequences and other useful features (all described below) were added to the annotation. Finally, detected transcript borders (TSSs and terminators) were used to refine coordinates of annotated features (e.g. alternative translational initiation sites).	Afterwards, the quality of genome-wide translational initiation site (TIS) calls was evaluated using ‘assess_TIS_annotation.py’ (14).

A total of 178 publications (15–192) was used for curation of the genome annotation.

**Classification of encoded proteins**

Protein sequences corresponding to all CDS features in the final annotation were determined from the corresponding coding sequences. Using InterProScan (193) all proteins were classified according to several databases: TIGRFAMs (194), Gene Ontology (GO, (195, 196)), KEGG (197), InterPro (IPR, (198)), and Protein family (Pfam, (199)).

**Mapping of RNA-seq data**

For the mapping of RNA-seq data, the first 1000 bps of the newly assembled reference genome were copied to the end of the genome, to enable mapping of sequencing reads across the boundaries of the sequence file. Subsequently, we used Bowtie 2 (8) to map Illumina sequencing data (described above) to the extended genome in the appropriate mode (single-end for 5’-enriched, paired-end for control data).

**Normalized start and end counts and complete coverage of sequenced fragments**

The start position of sequenced fragments was extracted from the sequence alignment map (SAM) produced by Bowtie 2. In case of paired-end sequencing, end points of sequenced fragments were also extracted. The positions were used to build strand-specific, single-nucleotide resolution frequency tables (start counts, end counts and coverage). For single-end data, coverage was calculated from the regions covered by the single reads. On the other hand, for paired-end data, the entire inferred fragment (i.e. including the region between mapping sites of mate reads) was used to calculate coverage. Counts of the last 1000 bps of the artificially extended genome were added to those of the first 1000 bps. To obtain normalized values, start counts, end counts and coverage were each divided by the summed genome-wide coverage, excluding positions within 30 nts of rRNA genes.

**Identification of putative coverage termination peaks**

The end count frequency tables for each strand were scanned with window sizes of 3-12 nucleotides and the evaluated window was marked as a candidate termination peak if it met a set of criteria. Firstly, the average end count per nucleotide should be more than 10-fold higher inside the window than in the 8 nucleotides upstream and 8 nucleotides downstream of the window. Secondly, the average number of terminated reads per nucleotide should be higher than 5. Thirdly, peaks were trimmed when the most outer nucleotides had an end frequency of either below 3 or below 1% of the maximum frequency within the window. Then, overlapping candidate peaks were merged and the merged peaks were trimmed as before. The final set of peaks was named ‘putative coverage termination peaks’, shown in PneumoBrowse.

**Detection of small RNA features**

Paired-end sequencing was exploited to detect putative small RNA features. To be conservative, only uniquely mapped read pairs were processed, effectively eliminating reads mapping to rRNA regions. This is the only instance where multi-mappers were excluded. First, to determine the distribution of randomly fragmented transcripts, the sizes of sequenced fragments were extracted from the SAM file corresponding to the paired-end-sequenced control sample. Herein, fragments (partially) mapping within 30 nts upstream of or overlapping with annotated features smaller than 500 nts were excluded. This was done to exclude biases towards lower fragment sizes. In addition, the 10 nts downstream of tRNA genes were excluded, to eliminate fragments derived from tRNA processing. From all remaining fragments, the library-wide size distribution was determined.

Subsequently, for each putative coverage termination peak (see above), fragments from the SAM file that ended inside the peak region were extracted. Of those reads, a peak-specific fragment size distribution was built. A putative sRNA was defined by several criteria: (i) the termination efficiency of the coverage termination peak should be above 30% (see below for the definition of termination efficiency), (ii) the relative frequency of the predicted sRNA length should be more than 25-fold higher than the corresponding frequency in the library-wide distribution, (iii) the predicted sRNA should be completely covered at least 15x for high-confidence (HC) terminators and at least 200x for non-HC terminators (see below for the definition of HC terminators).

The entire process was repeated once more, now also excluding all detected putative sRNAs from the library-wide size distribution. This allowed detection of lower-abundance sRNAs that were of the same size as a highly abundant sRNA identified in the first round. For the scope of this paper, only predicted sRNAs that did not significantly overlap already annotated features were considered.

A candidate sRNA was annotated (either as sRNA or riboswitch) when either (i) a matching entry, with a specified function, was found in RFAM (200) and/or BSRD (145) databases (e.g. T-box leaders, PyrR-binding sites); (ii) the sRNA was validated by Northern blotting, either in previous studies or by us; or (iii) at least two transcription-regulatory elements were detected (i.e. transcriptional start or termination sites, or sigma factor binding sites).

**Northern blotting**

Cells for Northern analysis were grown in C+Y medium. At OD_600_ ~ 0.1, 100 ng·μl^-1^ CSP-1 was added and cells were harvested 20 min afterwards. Total RNA was isolated and Northern blotting was performed as described by Van der Meulen et al. (van der Meulen et al. 2016), but using 2% bleach instead of 7M urea in the separating polyacrylamide gel. Probe sequences are listed in **Supplementary Table S2**.

**Identifying high-confidence terminators**

Since *S. pneumoniae* lacks the Rho factor, all terminators are expected to be ‘Rho-independent’ or intrinsic terminators. These intrinsic terminators consist of a palindromic stem, the arms of which are separated by a small loop, followed by a uracil-rich stretch. Using the new genome assembly as input, putative stem loops followed by U-rich sequences were predicted by TransTermHP (1), with liberal settings to reduce false negative rates: a minimum confidence level of 60, a maximum loop length of 15 (with a penalty increasing linearly with length) and no genomic context restrictions. Subsequently, if a previously identified ‘putative coverage termination peak’ fell inside the poly(U)-tract of a predicted stem-loop structure, the combination of both elements was defined to be a HC-terminator and was included in the genome annotation. Terminator efficiency was determined by the total number of fragments ending in a coverage termination peak, as a percentage of all fragments covering the peak (i.e. including non-terminated fragments).

**Transcription start site identification**

Normalized start counts from 5’-enriched and control libraries were compared. Importantly, normalization was performed excluding reads that mapped to rRNA genes, to account for the fact that the control library was and the 5’-enriched library was not rRNA-depleted. Additionally, all locations within 30 bps of rRNA genes were excluded from further analysis. An initial list was built of unclustered TSSs, which have (i) at least 2.5-fold higher normalized start counts in the 5’-enriched library, compared to the control library, and (ii) a minimum normalized start count of 2 (corresponding to 29 reads) in the 5’-enriched library. Subsequently, TSS candidates closer than 10 nucleotides were clustered, conserving the candidate with the highest start count in the 5’-enriched library. Finally, if the 5’-enriched start count of a candidate TSS was exceeded by the value at the nucleotide immediately upstream, the latter was annotated as TSS instead. The remaining, clustered TSSs are referred to as high-confidence (HC) TSSs.

The 5’-end of some transcripts may be rapidly dephosphorylated, precluding their enrichment in the Cappable-seq approach. Therefore, we included a set of 34 lower confidence (LC) TSSs in our annotation, which were not overrepresented in the 5’-enriched library, but that did meet a set of strict criteria: (i) normalized start count in the control library should be above 10 (corresponding to 222 reads), (ii) a TATAAT motif (with a maximum of 1 mismatch) should be present in the 5-15 nucleotides upstream, (iii) the nucleotide should not be immediately downstream of a processed tRNA, and (iv) the nucleotide should be in an intergenic region. If multiple LC-TSSs were predicted in one intergenic region, only the strongest one was annotated. If a HC-TSS was present in the same intergenic region, the LC-TSS was only annotated when its 5’-enriched start count exceeded that of the HC-TSS.

**TSS classification**

After transcript and CDS boundaries were refined, using the detected transcript boundaries (TSSs and terminators), the final set of high- and low-confidence TSSs were classified into five classes: primary TSSs (only or strongest TSS within 300 nt upstream of a feature), secondary TSSs (within 300 nt upstream of a feature, but not the strongest), internal (inside a feature and on the same strand), antisense (inside a feature or untranslated region, on the opposite strand), and orphan (in none of the other four classes). Antisense TSSs were further divided into 3 subclasses: A_5_ (antisense and upstream of feature), A_3_ (antisense and downstream of feature), and A_0_ (antisense and inside feature).

**Prediction of regulatory motifs**

We proceeded with the identification of binding sites of sigma and transcription factors. First we retrieved reported binding sites for RpoD (σ_A_, (187)), ComX (σ_X_, (67)) and ComE (146), along with pneumococcal extended -10 sequences (201). Using the MEME Suite (113), we built consensus binding motifs: the nucleotide distribution of all regions of 500 nucleotides upstream of TSSs was used as background nucleotide distribution. For RpoD, separate motifs were built for -35 and -10 sequences and five composite motifs were then created with a spacer length of 15-19 nucleotides between the -35 and -10 elements.

Subsequently, using FIMO (202), the 100 nucleotides upstream of each TSS were scanned for a match with the consensus ComE-binding motif (p<0.00001). Similarly, the 40 nucleotides upstream were searched for sites recognized by ComX (p<0.00001) or RpoD (p<0.001). Afterwards, predicted RpoD binding sites were discarded if the spacing between the motif and the corresponding TSS was below 3 or above 8. The spacing between a ComX site and a TSS was required to be below 6. Finally, a search for extended -10 sites was limited to 20 nucleotides upstream of TSSs (p<0.001).

Annotated binding sites of other transcription factors, including CodY and CcpA, were directly adopted from RegPrecise (157).

Ribosomal binding sites (RBSs) were identified by scanning the 16 nucleotides upstream of translation initiation sites (TISs) with MEME, identifying an enriched motif (**Figure 7A**, inset), present for 69% of TISs (p<0.01). Spacing between RBSs and corresponding TISs was restricted to 4-9 nucleotides.

The origin of replication was predicted with OriFinder (101). DnaA-boxes were annotated exclusively in the 5 kbp region surrounding the origin of replication. Perfect matches to the consensus sequence TTWTNCACA (28) were supplemented with single-mismatch sequences located within 5 nucleotides of a perfect match. DnaA-trios were annotated based on sequence identity with their *B. subtilis* counterparts (179).

**Operon prediction and regulon assignment**

Defining an operon as a set of genes controlled by a single promoter, putative operons were predicted for each primary TSS, as defined above. Two consecutive features on the same strand were predicted to be in the same operon if (i) their expression across 22 infection-relevant conditions was strongly correlated (correlation value > 0.75 (BioRxiv: <https://doi.org/10.1101/283739>)) and (ii) no strong terminator (>80% efficient) was found between the features. The resulting operons (**Supplementary Table S8**) were allowed to overlap, reflecting the complex nature of the transcriptome. Regulons were assigned according to detected transcription factor sites upstream of primary TSSs, combined with predicted operons.

**Identification of leaderless transcripts**

For all 767 operons starting with a protein-encoding feature (CDS) or pseudogene, the TSS position was compared to the start position of the first feature in the operon. Thereby, 5’-leaders were determined. We revealed 80 operons with a leader of 10 nucleotides or less, while 69 of them had no leader at all.

**Creating PneumoBrowse**

PneumoBrowse (<https://veeninglab.com/pneumobrowse>) is based on JBrowse (203), supplemented with plugins *jbrowse-dark-theme* and *SitewideNotices* (<https://github.com/erasche>), and *ScreenShotPlugin*, *HierarchicalCheckboxPlugin* and *StrandedPlotPlugin* (BioRxiv: <https://doi.org/10.1101/212654>). Annotated elements were divided over five annotation tracks: (i) genes (includes pseudogenes, shown in grey), (ii) putative operons, (iii) regulatory features, including TSSs and terminators, (iv) repeats, and (v) other features. Additionally, full coverage tracks are available, along with start and end counts.

9. Tatusova,T., DiCuccio,M., Badretdin,A., Chetvernin,V., Ciufo,S. and Li,W. (2013) Prokaryotic Genome Annotation Pipeline. In *The NCBI Handbook*. National Center for Biotechnology Information (US).
